# Unilateral auditory deprivation reveals brainstem origin of a sensitive period for spatial hearing

**DOI:** 10.1101/2024.04.01.587638

**Authors:** Kelsey L. Anbuhl, Alexander T. Ferber, Andrew D. Brown, Victor Benichoux, Nathaniel T. Greene, Daniel J. Tollin

## Abstract

Early sensory experience can exert lasting perceptual consequences. For example, a brief period of auditory deprivation early in life can lead to persistent spatial hearing deficits. Some forms of hearing loss (i.e., conductive; CHL) can distort acoustical cues needed for spatial hearing, which depend on inputs from both ears. We hypothesize that asymmetric acoustic input during development disrupts auditory circuits that integrate binaural information. Here, we identify prolonged maturation of the binaural auditory brainstem in the guinea pig by tracking auditory evoked potentials across development. Using this age range, we induce a reversible unilateral CHL and ask whether behavioral and neural maturation are disrupted. We find that developmental CHL alters a brainstem readout of binaural function which is not observed when the CHL is induced in adulthood. Startle-based behavioral measures reveal poorer spatial resolution of sound sources, but only for high-frequency sound stimuli. Finally, single-unit recordings of auditory midbrain neurons reveal significantly poorer neural acuity to a sound location cue that largely depends on high-frequency sounds. Thus, these findings show that unilateral deprivation can disrupt developing auditory circuits that integrate binaural information and may give rise to lingering spatial hearing deficits.

## Introduction

Asymmetric sensory deprivation in development can have profound long-lasting effects on sensory systems that require bilateral integration of sensory input. Classically, monocular visual deprivation or eyelid closure during discrete developmental periods results in distinct structural and functional changes to primary visual cortex neurons (Wiesel and Hubel, 1963; Hubel and Wiesel, 1970; Maffei et al., 2004; Hofer et al., 2009; Coleman et al., 2010; Espinosa and Stryker, 2012; Zhou et al., 2017), along with deficits in visually-guided behavior that require binocular perception (Dews and Wiesel, 1970; Blake and Hirsch, 1975; Van Hof-Van Duin, 1976; Timney et al., 1980). Unilateral whisker trimming or other mechanical manipulation alters dendritic morphology and synaptic physiology of primary somatosensory cortex neurons, accompanied with degraded neural response properties in the somatotopically-organized barrel cortex (Fox, 1992; Vees et al., 1999; Allen et al., 2003; Rema et al., 2003; Shepherd et al., 2003; Bender, 2006). Similarly, auditory deprivation in one ear can also disrupt the response properties of neurons in the auditory cortex (Brugge et al., 1985; Popescu and Polley, 2010; Keating et al., 2013a, 2015; Polley et al., 2013), as well as alter the morphology, connectivity, and response properties of other central auditory nuclei (Kumpik and King, 2018). Thus, central circuits that require coordinated integration of bilateral sensory inputs appear to be particularly vulnerable to unilateral deprivation during developmental sensitive periods. In fact, brief periods of reversible unilateral hearing loss in developing animals can disrupt binaural selectivity of central auditory neurons in the auditory midbrain (Clopton and Silverman, 1977; Silverman and Clopton, 1977; Mogdans and Knudsen, 1993; Xu and Jen, 2001; Popescu and Polley, 2010), thalamus (Miller and Knudsen, 2003), and cortex (Brugge et al., 1985; Popescu and Polley, 2010; Keating et al., 2013a, 2015; Polley et al., 2013) with concurrent disruptions to binaural and spatial hearing abilities (*for review*: Gordon & Kral, 2019; Kumpik & King, 2019). It is not clear, however, whether early unilateral deprivation can also induce plasticity at the auditory brainstem, the initial site of synaptic convergence between input from both ears (Kandler et al., 2009).

The superior olivary complex in the auditory brainstem contains neural circuits specialized to extract subtle differences between the times of arrival (interaural time difference, ITDs) and amplitudes (interaural level difference, ILDs) of sound stimuli at both ears (Owrutsky et al., 2021; Tollin, 2003). The binaural cues to sound location, ITDs and ILDs, allow for the localization of sound in the horizontal plane and are initially computed at the medial and lateral superior olive (MSO, LSO), respectively. Neurons in the LSO encode ILDs and onset ITDs by integrating excitatory, glutamatergic inputs from the ipsilateral cochlear nucleus with inhibitory, glycinergic inputs from the contralateral ear via large synapses from the medial nucleus of the trapezoid body (MNTB) (Sanes, 1990a; Tollin, 2003). After hearing onset, the superior olivary cochlear nuclei display significant synaptic reorganization of tonotopic maps which is not observed in the cochlear nucleus, a monaural structure (Kandler et al., 2009). This refinement of tonotopy is likely to reflect experience-dependent alignment of frequency-specific inputs from each ear, which is evident in the pruning of MNTB axon terminals and LSO dendrites and the improvement of frequency selectivity in developing gerbils after hearing onset (Sanes and Rubel, 1988). This suggests that binaural circuits might be vulnerable to asymmetrical hearing loss, especially considering neurons in the LSO integrate ipsilateral and contralateral afferents that need to be temporally and spectrally-matched to maintain precision in ILD and ITD encoding. Indeed, unilateral deafferentation of the cochlea disrupts specificity of MNTB arborizations (Sanes and Takács, 1993).

Children that experience conductive hearing loss (CHL) due to otitis media with effusion (i.e., accumulation of fluid in the middle ear) often display binaural hearing impairments that persist months to years after resolution of the hearing loss and restoration of normal audibility (Bennett et al., 2001; Hall III and Grose, 1994; Hall et al., 1995; Ludwig et al., 2019; Moore et al., 2003; Reichman and Healey, 1983; Whitton and Polley, 2011). A physiologically-derived biomarker for binaural function− the binaural interaction component (BIC) of the auditory brainstem response (ABR)− has been shown to be significantly altered in children with a history of CHL (Gunnarson and Finitzo, 1991; Hall and Grose, 1993; Delb et al., 2003; Laumen et al., 2016). Given that the BIC has origins in the auditory brainstem (i.e., LSO; Ungan et al., 1997; Benichoux et al., 2018; Brown et al., 2019; Tolnai and Klump, 2020), it raises the possibility that early hearing loss can induce plasticity in the auditory brainstem.

To address this question, we first characterized the development of binaural function in a precocial rodent, the guinea pig, and determined whether unilateral hearing loss during this time perturbs the maturation of neural tuning of binaural information in the brainstem and midbrain. We found that a brainstem readout of binaural function (i.e., BIC of the ABR) displays a prolonged maturation period suggesting that binaural plasticity may be heightened during this time, and therefore vulnerable to auditory experience. Indeed, we found that a reversible unilateral hearing loss in development disrupts the BIC of the ABR, whereas hearing loss of the same duration induced in adult animals did not give rise to BIC impairments. We then asked whether early CHL also contributes to spatial hearing deficits. We found a significant reduction in spatial acuity (via startle-based methods) with high-pass, but not broadband, noise stimuli. Finally, *in vivo* single-unit recordings of auditory midbrain neurons revealed diminished neural acuity to interaural level difference (ILD) cues to sound location. Taken together, these results suggest that asymmetric auditory input during development can disrupt binaural circuits at the level of brainstem and midbrain that support spatial hearing, and may, in part, explain the spatial hearing deficits observed in animals and children following developmental CHL.

## Results

### Prolonged maturation of binaural auditory brainstem function

First, we sought to characterize the developmental time-course of binaural brainstem function and subsequently, how asymmetric perturbations of acoustic experience can alter this maturation. Studies examining binaural physiology have often used invasive techniques that prohibit the tracking of binaural neuron function within the same animal over time (Sanes and Rubel, 1988; Sanes, 1990b; Kim and Kandler, 2010). To circumvent these limitations, we used the auditory brainstem response (ABR) as a noninvasive measure of brainstem function. ABRs consist of a series of robust peaks representative of synchronous activation of neural structures along the ascending auditory pathway (Figure 1a). Here, we tracked ABR development in the guinea pig, a precocial mammal that shares similar auditory sensitivity as humans (Heffner et al., 1971; Prosen et al., 1978). Since the acoustical properties of the developing guinea stabilize and reach adult-like ranges around postnatal (P) day 56 (Anbuhl et al., 2017), we collected ABRs from birth (P1) through ages well beyond P56 (>P91) in a group of guinea pigs (n=18).

**Figure 1.**
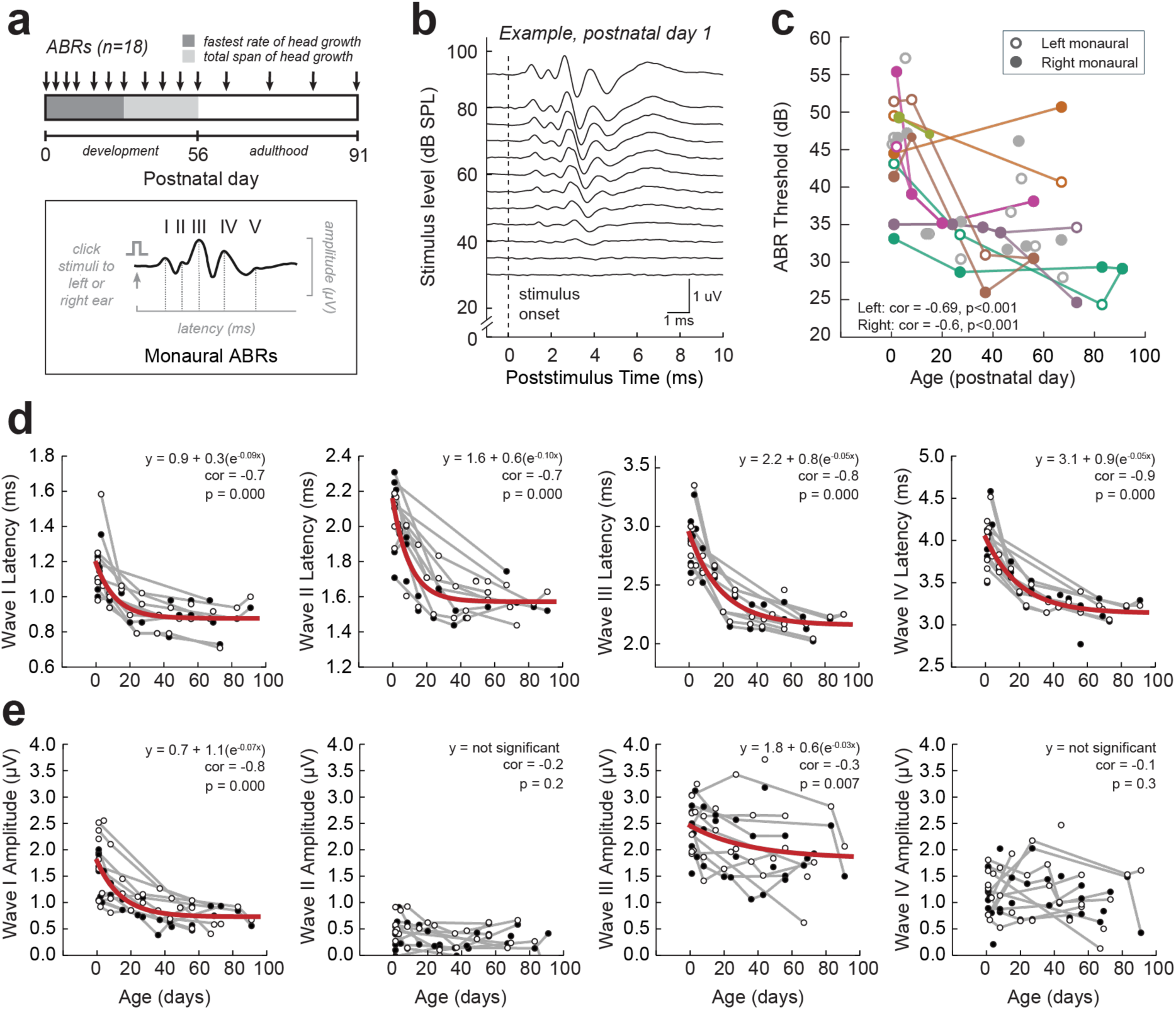
Development of guinea pig monaural auditory brainstem responses (ABRs). (**a**) *Inset*: The ABR is an auditory-evoked potential that displays peaks (black trace, waves I-IV) corresponding to locations along the ascending auditory pathway. Presentation of click stimuli to the left or right ear alone elicits a monaural ABR. *Timeline*: ABRs were collected in guinea pigs (n=18) at developmental ages spanning newborn (P1) through adulthood (>P56). A subset of guinea pigs (n=7) had ABR measurements tracked across development, and the remaining animals (n=11) had 1-2 ABRs at different developmental ages. (**b**) Example monaural ABR trace (right ear) for subject ID #124111 at postnatal (P) day 1. Broadband click stimuli are presented across a range of stimulus intensities (30-90 dB SPL in 5 dB steps). Traces shown are the average of 500 repetitions. The dotted vertical line indicates the stimulus onset of the click. Threshold is defined as the softest intensity click presented in order to elicit a distinctive ABR waveform (e.g., threshold for this example would be 40 dB SPL). (**c**) Click-ABR thresholds as a function of developmental age (P1-P90). Circles indicate individual thresholds from the left (open circles) and right ear (filled circles) ABR measurements. Lines connect data from the same animal tracked over time. Click-ABR thresholds decrease significantly with age (Left correlation = −0.69, p<0.001; Right correlation: −0.6, p<0.001). (**d, e**) Monaural ABR wave I-IV latency (**d**) and amplitude (**e**) plotted as a function of age. Each column indicates a different wave (I-IV). Circles indicate values from the left (open circle) or right ear (filled circle) monaural ABRs. Gray lines indicate data points corresponding to the same animal tracked over time. When a significant correlation is found, an exponential decay function is fit to the data (red line). Significant correlations are found for the amplitudes of monaural waves I and III (p=0.000-0.007) and latencies of waves I-IV (p=0.000 for all waves).

We first characterized the maturation of monaural ABRs. Figure 1b shows an example ABR from a newborn guinea pig (P1), where monaural clicks (i.e., presented to one ear alone) were presented across a range of sound intensities (30-90 dB SPL). ABR thresholds were defined as the softest intensity that generates a visible ABR waveform. We find that monaural ABR thresholds decreased significantly with age (*Spearman’s rho correlation*: Left monaural, r_s_ = −0.71, n = 38, p = 0.000; Right monaural, r_s_ = −0.54, n = 38, p = 0.002), with the greatest decrease occurring in the first 1-2 postnatal weeks (Figure 1c). We then examined individual ABR waves for developmental changes in wave amplitude and latency, which are reflective of synchronous sound-evoked neural activity and timing of the ascending auditory nuclei, respectively. The guinea pig exhibits four to five distinct ABR waves (Dum, 1984; Figure 1a), where waves I-III are generated by activation of the auditory nerve and cochlear nucleus, and waves IV-V are activated by the superior olivary complex and the lateral lemniscal tract leading to the auditory midbrain (i.e., inferior colliculus). We find that the amplitudes of monaural waves I and III significantly decrease with age (wave I: r_s_ = −0.75, p = 0.000; wave II: r_s_ = −0.16, p = 0.223; wave III: r_s_ = −0.34, p = 0.007; wave IV: r_s_ = −0.13, p = 0.331), along with a significant decrease in latencies of all monaural waves (wave I: r_s_ = −0.72, p = 0.000; wave II: r_s_ = −0.73, p = 0.000; wave III: r_s_ = −0.83, p = 0.000; wave IV: r_s_ = −0.85, p = 0.000; Figure 1d, e). In general, the latency of the earlier waves stabilized by ∼P21 and the later waves stabilized by ∼P56 suggesting that more peripheral monaural auditory structures (i.e., auditory nerve, cochlear nucleus) mature earlier in postnatal development than later peaks associated with superior olivary complex nuclei.

We next characterized binaural function using the ABR measurements. Since the ABR includes evoked responses from the superior olivary complex (medial and lateral superior olives; MSO, LSO; Figure 2a), where the initial processing of the binaural acoustical cues occurs (Owrutsky et al., 2021), it is possible to derive a binaural-specific component that is separate from the monaural response. Due to the monaural origin of the early ABR waves, the magnitude of the peaks observed in response to binaural clicks (i.e., presented to both ears) is equal to the summed magnitudes in response to monaural clicks (Figure 2c). Later peaks that are binaural in origin are not equal to the sum of their monaural ABRs, indicating a binaural-specific computation (Jewett, 1970). This difference is referred to as the binaural interaction component (BIC; Figure 2c). We focused on the largest component of the BIC, called DN1, which is altered in patients with binaural hearing dysfunction despite having normal audiological hearing (Laumen et al., 2016).

**Figure 2.**
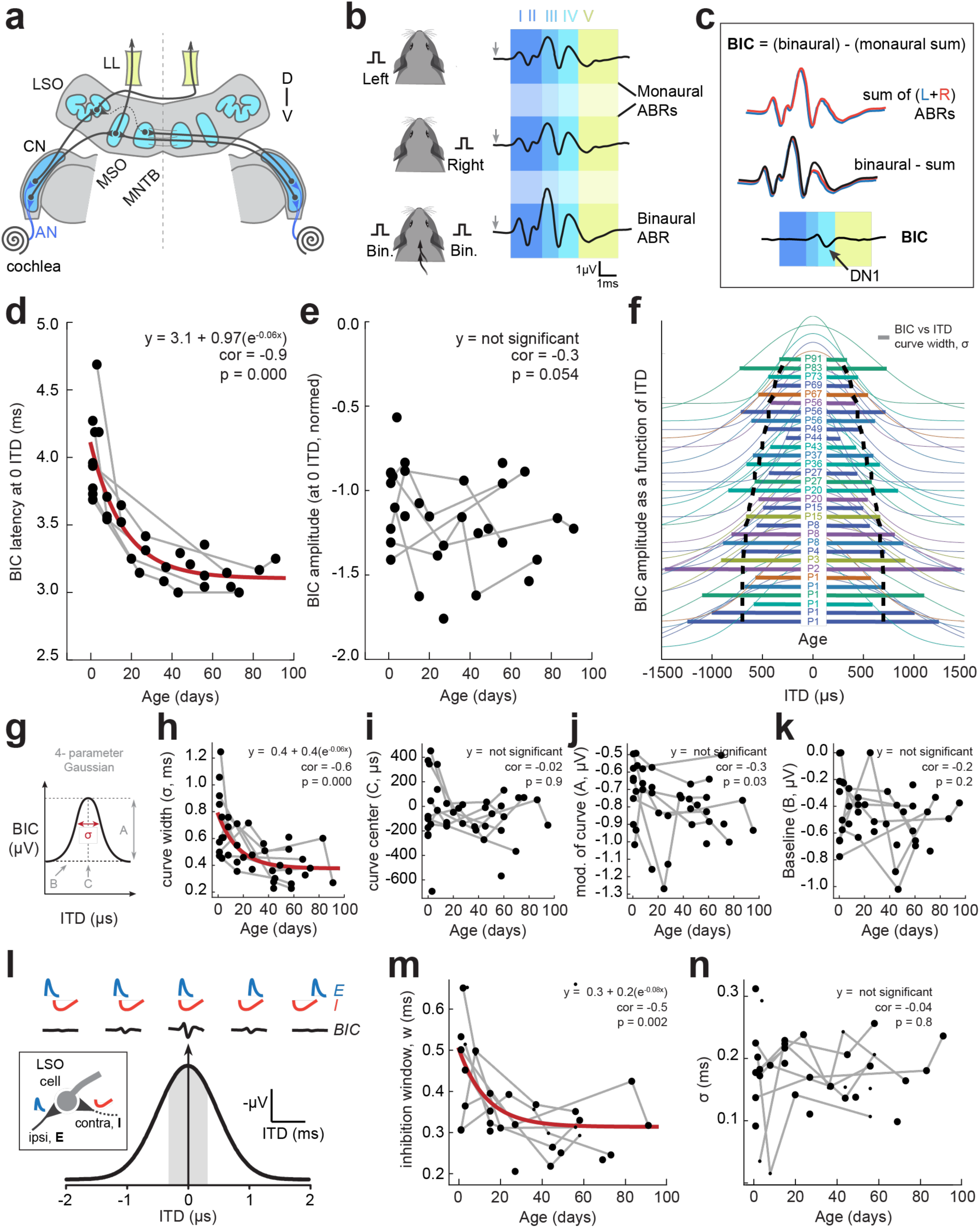
Development of binaural auditory brainstem physiology. (**a**) Cross-section through the auditory brainstem and the neural generators of the auditory brainstem response (ABR): AN=auditory nerve, CN=cochlear nucleus, LL=lateral lemniscus, LSO=lateral superior olive, MSO=medial superior olive, MNTB=medial nucleus of the trapezoid body. D=dorsal, V=ventral. (**b**) The ABR is an auditory-evoked potential that results in peaks (black trace) corresponding to locations along the ascending auditory pathway. Presentation of click stimuli to the left ear or right ear alone elicits a monaural ABR. Presentation of click stimuli to both ears elicits a binaural ABR. (**c**) When the left and right monaural ABRs are summed and compared to the binaural ABR, there is a difference in potentials in ABR wave IV, which corresponds to the superior olivary complex (i.e., LSO, MSO) a location that integrates binaural information. The difference in binaural and the summed monaural potential gives rise to the binaural interaction component (BIC), which contains the DN1 peak (see arrow). (**d**) ABRs were collected in guinea pigs (n=18) at developmental ages spanning newborn (∼P1) through adulthood (>P56). A subset of guinea pigs (n=7) had ABR measurements tracked across development, and the remaining animals (n=11) had 1-2 ABRs at different developmental ages. The BIC was computed for each ABR assessment. The latency of the BIC (at 0 µs ITD) decreased with age. The Spearman’s rho correlation analysis revealed this decrease to be significant (p=0.000), and an exponential decay function was fit to the data (red line). (**e**) Normalized amplitude of the BIC (at 0 µs ITD) was not significantly effected by age (Spearman’s rho correlation, p=0.054). Gray lines: data points from the same animal tracked over time. (**f**) BIC amplitude as a function of ITD (fitted with a gaussian model; centered and normalized) for each ABR timepoint across development. Colors indicate each animal tested. Horizontal lines indicate the curve width (σ) of each fitted function. The functions are stacked in descending age order (age listed in center). The black dotted line is the regression exponential fitted to the curve widths, and show a decrease in BIC vs ITD curve width with age. (**g-k**) Development of BIC amplitude versus ITD fitted functions. (**g**) BIC amplitude vs ITD data were fit with a 4-parameter Gaussian (σ, the curve width; C, the curve center; A, the modulation of the curve; B, the baseline value). The Spearman’s rho correlation analysis revealed a significant decrease in the curve width of the function with age (**h**; correlation= −0.63, p=0.000) but not for the other three parameters (**i-k**; correlations = −0.34 to −0.02, p=0.032-0.901). An exponential decay function was fit to the curve width data (h; red line). (**l**) The BIC likely derives from an excitatory (E)-inhibitory (I) computation present in the LSO. BIC amplitude is maximum at 0 µs ITD, which corresponds to the greatest E-I overlap (i.e., largest reduction in the binaural waveform). The grey shaded area corresponds to the acoustical ITD range for the guinea pig. (**m-o**) A model of binaural interaction of LSO cells was used with two parameters: sigma (σ), representing the precision of arrival times of excitatory and inhibitory inputs to LSO cells, and the inhibition window (w), representing the temporal duration of the inhibition. The window of inhibition (w) decreased significantly with age (**m**; correlation= −0.49; p=0.002), whereas the σ parameter did not (**n**; p=0.835), An exponential decay function (red line) was fit if a significant correlation was found. All R2 values of the fits were >0.7 with a majority of fits >0.9.

The derived binaural-specific BIC of the ABR displayed a significant decrease in latency with age (r_s_ = −0.85, n = 39, p = 0.000), similar to the developmental time course of the later ABR waves (Figure 2d). No significant change in BIC amplitude with age was observed (p = 0.054; Figure 2e). Since the BIC is derived by the excitatory-inhibitory (EI) interaction within the LSO of the auditory brainstem (Ungan et al., 1997; Benichoux et al., 2018; Brown et al., 2019; Tolnai and Klump, 2020), the timing of click presentations to each ear can be leveraged as a tool for examining the maturation of EI interactions. By varying the interaural timing of clicks to both ears (i.e., the interaural time difference, ITD), we can generate a bell-shaped BIC amplitude versus ITD curve where amplitude is maximal for 0 μs ITDs (i.e., no temporal disparity between left and right ear clicks) and tapers off with non-zero ITDs (Ferber et al., 2016b) (Figure 2f). The BIC vs ITD data are fit with a 4-parameter Gaussian function where we quantify: (1) the ITD of maximal BIC, (2) the curve width (σ) of the Gaussian function, (3) the modulation of the curve (i.e., amplitude), and (4) the baseline BIC value (Figure 2g). Spearman’s rho correlation analysis revealed a significant decrease in the curve width with age (r_s_ = −0.63, p = 0.000), but not for the other three parameters (correlations: −0.34 to −0.02, p = 0.032-0.901). (Figure 2f,h;). Figure 2f shows individual BIC vs ITD functions for each timepoint collected, along with horizontal lines showing the decreasing curve width (σ) with age (black dotted line). Newborn animals (P1) exhibit broad BIC vs ITD curves. As animals age, curve widths narrow and stabilize after ∼P40.

To determine the source of the narrowing of BIC vs. ITD curve width with age, we used a previously established model (Benichoux et al., 2018) to pinpoint whether: (1) the arrival time and resultant interaction of excitatory and inhibitory inputs to the LSO becomes more precise with arrival times of EI inputs, and/or (2) the window of EI integration shortens as the auditory brainstem matures (Figure 2l). The model fit the data well (R^2^ 0.7-0.99) and revealed the EI integration window in LSO cells significantly decreased with age from ∼0.5 ms in newborn animals to ∼0.3 ms at ages >P40 (Figure 2m; r_s_ = −0.49, p = 0.002). Age had no significant effect on the precision of EI arrival time (r_s_ = −0.04, p = 0.835; Figure 2n).

In summary, ABRs assessments across age revealed the maturation of the guinea pig auditory brainstem follows a similar developmental time-course as the acoustical cues to sound location (Anbuhl et al., 2017), where values are stable by P56. Therefore, this provides the precise duration needed to induce a unilateral hearing loss that spans binaural hearing development.

### Early unilateral hearing loss alters the binaural interaction component of the ABR

Since the auditory brainstem of the guinea pig matures over a prolonged period, this suggests that the binaural circuits may be vulnerable to sensory deprivation during this time. Thus, we hypothesized that a hearing loss in one ear would disrupt the normal development of binaural circuits that depend on input from both ears. To test this, we induced a reversible, mild-moderate hearing loss in one ear using custom earplugs (Figure 3a,b) from P0 (birth) through adulthood (P56) in guinea pigs. When animals reached P56, the earplug was removed, and normal auditory input was restored (see timeline in Figure 3c). The earplug manipulation was used as it allows for a conductive hearing loss (CHL) that is reversible and without permanent disruption of cochlear function. Animals that received a developmental CHL (n=21) were designated as “Early CHL” and compared to littermate controls that did not receive an earplug (“Control”; n=38).

**Figure 3.**
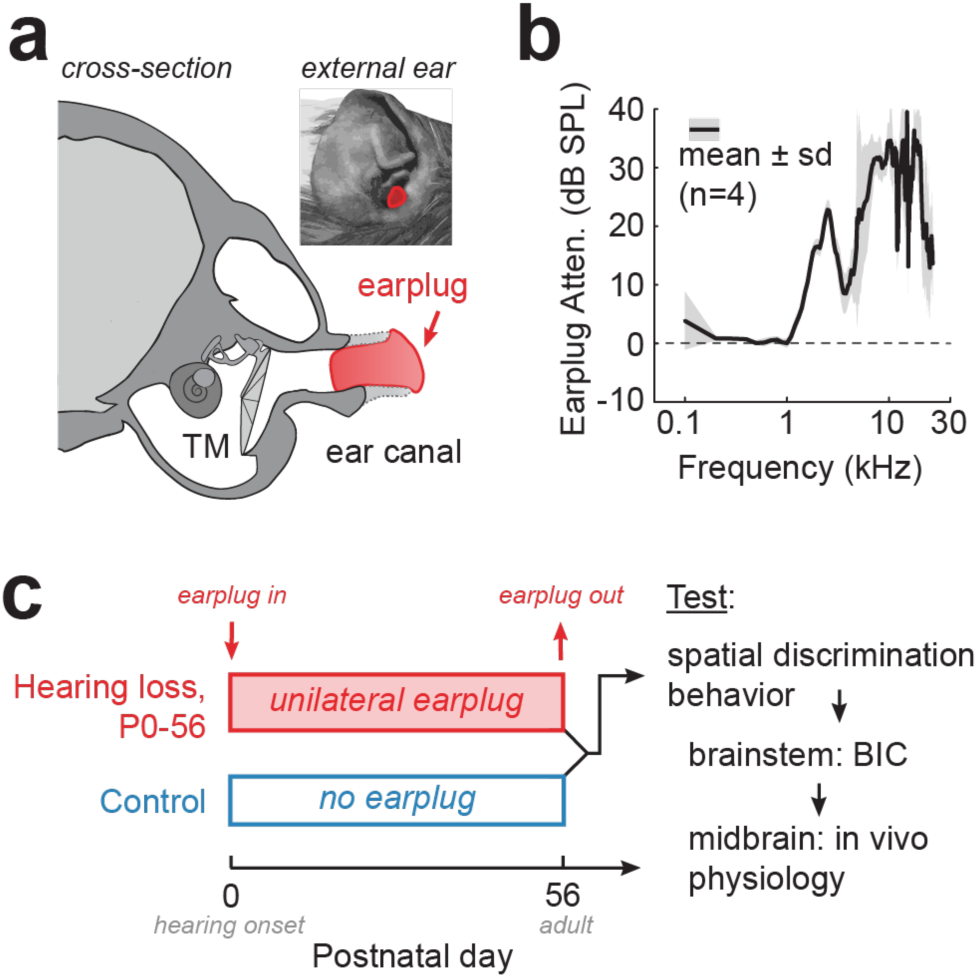
Transient conductive hearing loss (CHL) during development. (**a**) Schematic of coronal section of a guinea pig skull (traced from high-resolution CT scan). A custom-made earplug is fitted within the cartilaginous ear canal (dotted line), a safe distance from the tympanic membrane (TM). Inset: View of external ear with earplug in place. (**b**) Earplugs provide 10-35 dB sound attenuation for frequencies greater than 1kHz. (**c**) Experimental timeline. Litters of newborn guinea pigs were divided into two groups at birth: pups that were raised with no earplug (littermate Controls) and pups that were raised with a unilateral earplug (”early CHL”). Earplugs remained in place until adulthood (P56). Following earplug removal, animals underwent the following (in this order): a startle-based spatial discrimination task, auditory brainstem responses (ABRs) including the binaural interaction component (BIC), and in vivo single unit recordings in the auditory midbrain (i.e., inferior colliculus, IC).

After earplugs were removed at P56, ABRs were obtained in Early CHL and Control animals. First, we verified whether normal audibility was restored following earplug removal. We examined click-evoked monaural ABRs and determined ABR thresholds for each animal (see example in Supplemental Figure 1a). For Early CHL animals, monaural ABRs from the ear ipsilateral to the previous CHL are designated as “Early CHL_ipsi_” and “Early CHL_contra_” for monaural ABRs from the ear contralateral to the previous CHL. Indeed, both normal hearing and CHL-reared animals displayed comparable click-evoked monaural ABR thresholds (median ± SD: Control: 35±7.2 dB SPL; Early CHL_ipsi_: 40±7.9 dB SPL; Early CHL_contra_: 35±5.8 dB SPL; Mann-Whitney U: p>0.01; Supplemental Figure 1b). Developmental CHL did not significantly alter ABR wave I latencies compared to controls (Mann-Whitney U: p>0.05; Supplemental Figure 1c). The latencies for ABR waves III and IV were comparable between control and Early CHL_contra_ (Mann-Whitney U: p>0.05; Supplemental Figure 1d,e), though Early CHL_ipsi_ displayed longer latencies for monaural ABR waves III and IV (Mann-Whitney U; wave III: p<0.05; wave IV: p<0.01; Supplemental Figure 1d,e). Developmental CHL also did not significantly alter monaural ABR wave I-IV amplitudes compared to controls (Mann-Whitney U: p>0.05; Supplemental Figure 1f-h).

Since hearing loss has been reported to alter the latency of auditory brainstem responses of binaural nuclei (Folsom et al., 1983; Gunnarson and Finitzo, 1991; Laska et al., 1992; Hurley and Hurley, 1995), we first opted to examine BIC latency in Control and Early CHL animals. We find the median (±SD) latency for controls to be 3.25±0.19 ms, and for Early CHL to be 3.38±0.18 ms (Supplemental Figure 2). A Mann-Whitney U test revealed these latencies to be significantly different from one another (z=2.13, p=0.033), with Early CHL animals displaying, on average, 0.13 ms longer BIC latencies than Controls.

Next, we examined BIC tuning to interaural timing disparities (i.e., ITD) to probe underlying excitatory/inhibitory function of the auditory brainstem (Figure 4a, b). Figure 4c shows BIC amplitude versus the timing difference click stimuli presented to both ears (±0, 125, 250, 375, 500, 750, 1000, and 2000 μs) for Control (n=19 animals) and Early CHL animals (n=21). For both groups, the amplitude is maximal for an ITD of zero (i.e., no timing difference between ears) and decreases with larger ITDs (i.e., larger timing differences between ears). We fit the data with 4-parameter Gaussian functions (Figure 4b) that allowed us to quantify four parameters of BIC vs ITD functions between Controls and Early CHL animals: the ITD of maximal BIC, the curve width of the Gaussian function, the modulation of the curve (e.g., amplitude), and the baseline BIC value. The average BIC amplitude (±SEM) was computed at each ITD for controls and early CHL measurements and a Gaussian function was fit to each (Figure 4c). Of the four parameters, there is a significant difference between Controls and Early CHL for parameters A (modulation of the curve, z=-2.00, p=0.046; Figure 4d) and σ (curve width, z=2.95, p=0.0032; Figure 4e), but not for parameters B (baseline, z=-1.84, p=0.066; Figure 4f) or C (curve center, z=-0.79, p=0.43; Figure 4g). In summary, ABR measurements from Early CHL animals had significantly reduced BIC amplitudes at small ITDs (<250 μs) and significantly broader BIC vs ITD tuning compared to their littermate controls. These results suggest a unilateral CHL in development alters the maturation of binaural information processing at the level of the auditory brainstem.

**Figure 4.**
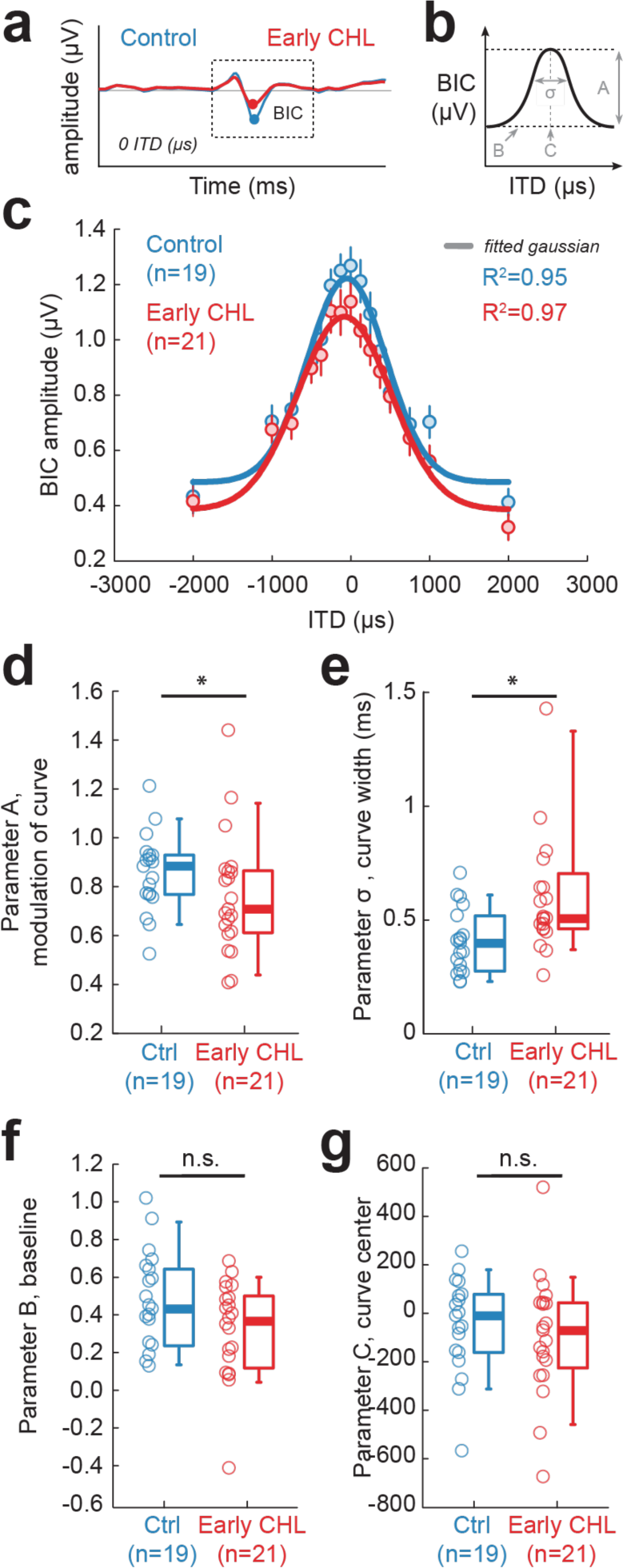
Unilateral developmental CHL alters the binaural interaction component of the ABR. (**a**) Example Control (blue) and Early CHL (red) binaural interaction component (BIC) traces at 0 ITD (µs). (**b**) BIC amplitude vs ITD data are fit with a 4-parameter Gaussian model. The parameters include: Parameter A, the modulation of the curve; Parameter B, baseline; Parameter C, the curve center; Parameter σ, curve width. (**c**) The average BIC amplitudes (µV; circles±SEM) were computed at each ITD tested (± 2ms) for control (n=19) and Early CHL animals (n=21). Gaussian functions were fit to the average BIC amplitude vs ITD for each group (thick lines). (**d**) Early CHL animals display significantly shallower modulation of the fitted gaussian curves (Parameter A) compared to Controls (Mann-Whitney U, p<0.05). (**e**) Early CHL animals display significantly broader gaussian curve widths compared to controls (curve width, σ) compared to Controls (Mann-Whitney U, p<0.05). (**f, g**) There were no significant differences between Early CHL and Controls for Parameter B (baseline) or Parameter C (curve center). Open circles indicate individual data, and boxplots indicate group medians.

### Adult-onset CHL does not disrupt the binaural interaction component of the ABR

Although we observed BIC alterations following 8 weeks of Early CHL, it is possible that the duration of CHL alone, at any age range, could lead to the same outcome. To test this, we induced unilateral CHL of the same duration as the Early CHL group (8 weeks) in adult guinea pigs (>P56; n=4) and measured BIC vs ITD tuning. Adults served as within-animal controls, where ABRs were collected prior to earplug placement (“Pre-CHL”; 3 sessions per animal) to establish baseline. Earplugs were then placed for 8 weeks, removed, and post-CHL ABR assessments were made (“Post-CHL”; 2 sessions per animal; see timeline in Figure 5a). Figure 5b shows the BIC amplitude (µV) as a function of ITD (µs) for Pre- and Post-CHL assessments. A 4-parameter Gaussian was fit to the average BIC amplitude for each ITD tested. We find that Adult-onset CHL did not significantly alter Parameter A (modulation of the curve; p=0.97; Figure 5c), curve width (p=0.97; Figure 5d), or Parameter B (baseline; p=0.46; Figure 5e) but did shift the curve center of the BIC vs ITD function (p=0.04; Figure 5f). This contrasts with the significant shifts observed following Early CHL with Parameter A (modulation of curve) and the curve width. Thus, we attribute the BIC alterations observed following Early CHL to be primarily restricted to developmental vulnerability to CHL.

**Figure 5.**
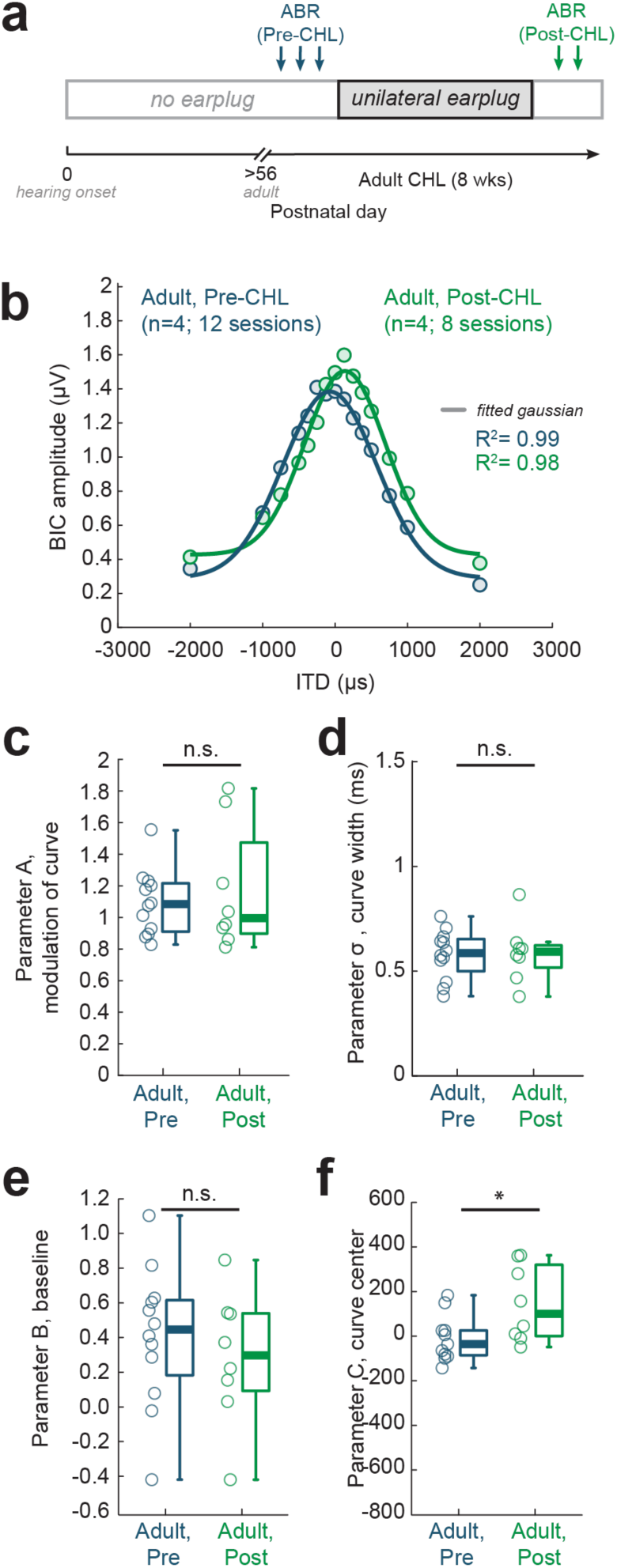
Adult-onset CHL has minimal impact on the binaural interaction component of the ABR. (**a**) Experimental timeline. ABRs were collected from normal-hearing Adult guinea pigs (>P56) prior to earplug placement (”Pre-CHL”; three sessions per animal). Unilateral earplugs were then placed in each animal for the same duration used in the developmental manipulation (8 weeks). Following earplug removal, ABRs were collected (”Post-CHL”; two sessions per animal). (**b**) BIC amplitude (µV) vs ITD (µs) for Pre-CHL and post-CHL ABR measures. The average for each condition (circles) are fit with a 4-parameter Gaussian model (thick line). (**c-f**) Adult-onset CHL did not alter Parameter A (modulation of the curve; p=0.97), curve width (p=0.97), or Parameter B (baseline; p=0.46), but did shift the BIC vs ITD curve center (p=0.04). Open circles indicate individual data, and boxplots indicate group medians.

### Early unilateral hearing loss disrupts spatial discrimination of high- but not low-frequency sounds

Since the binaural component of the ABR (i.e., BIC) was altered by early CHL, we asked whether Early CHL animals also displayed impairments in auditory behavior that requires binaural hearing. Previously employed behavioral assays are not optimal for our developmental study, as they use operant conditioning or other methods that require training that can take months (Heffner et al., 1971; Carney et al., 2011; Keating et al., 2013b) and are difficult to employ in some rodents including guinea pigs (Jonson et al., 1975). Therefore, we opted for an alternative approach that requires no training: a startle reflex-based method that can be used to assess spatial hearing ability in guinea pigs (Greene et al., 2018).

Behavioral spatial acuity was assessed using a speaker-swap paradigm (Allen and Ison, 2010; Greene et al., 2018; Figure 6a), which allowed us to determine the smallest angle for which a change in speaker location could be detected by the animal. High-pass or broadband noise was used to probe stimulus detection using acoustical cues available to the guinea pig. In the horizontal plane, broadband noise provides access to both interaural time and level difference cues (ITD, ILD), whereas high-pass noise largely provides access to ILD cues only. Guinea pigs reflexively startle in response to loud, unexpected sounds (Figure 6b), but presentation of a detectable cue (a “prepulse”) prior to the loud sound leads to a reduced startle response (Figure 6c). This is known as prepulse inhibition (PPI) of the acoustic startle response (Koch, 1999). Here, the prepulse is a shift of continuous noise stimulus (broadband or high-pass) presented from one speaker to a second speaker for a short interstimulus interval (ISI, 300 ms) prior to presentation of a startle-eliciting stimulus (via speaker mounted above animal). The direction of the swap was randomized (e.g. noise swaps from *left-to-right* or *right-to-left* speakers).

**Figure 6.**
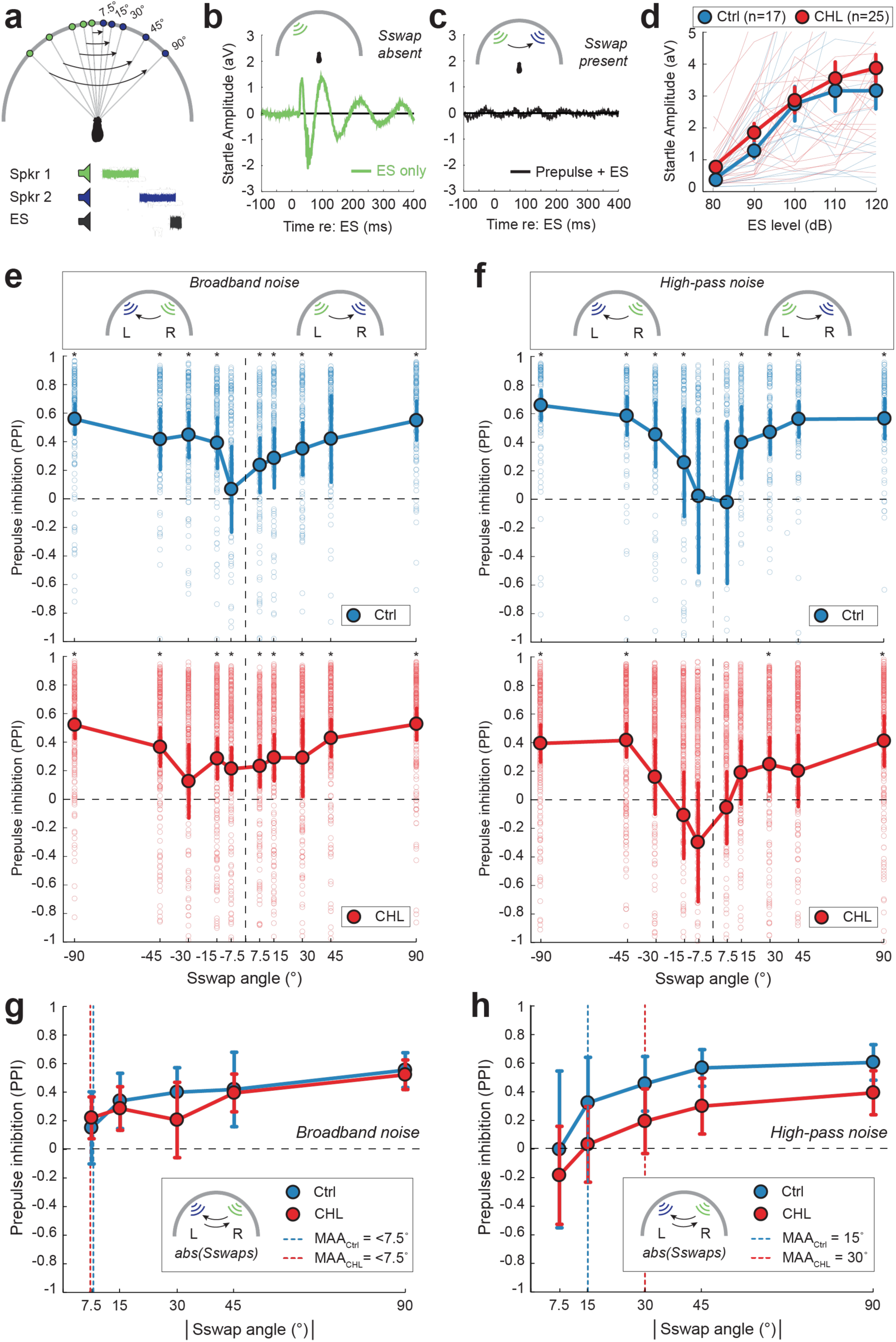
Unilateral developmental CHL alters spatial discrimination of high-pass noise. (**a**) Schematic of the startle-based behavioral paradigm to assess spatial discrimination. A continuous noise stimulus (low-pass or high-pass noise) is presented from one speaker (Spkr 1, green) then switches to a second speaker (Spkr 2, blue) for a short interstimulus interval (ISI, 300 ms) prior to presentation of a brief startle-eliciting stimulus (ES; 110-120 dB, 20 ms). The direction of the speaker swap (”Sswap”) is randomized (i.e., left-to-right, right-to-left). The angle of the speaker swap prepulse varied from 7.5° to 90° (7.5°, 15°, 30°, 45°, 90°). (**b**) Example startle response for the control condition where no prepulse was presented (ES only, no speaker swap). (**c**) Example response when a prepulse is presented prior to ES (Prepulse + ES). In this example, the animal detected the change in speaker locations resulting in a reduced startle response (Prepulse inhibition, PPI). (**d**) Startle response amplitude (arbitrary volts, aV) as a function of ES level (dB) in the presence of background noise (70 dB) for Control (n=17) and CHL (n=25) animals. There are no significant differences between groups in baseline startle responses (p>0.05). (**e**) Prepulse inhibition (PPI) as a function of speaker swap angle of broadband noise (100 Hz-20 kHz) for Controls (top panel, blue) and CHL animals (bottom panel, red). Data are shown for both right-to-left and left-to-right speaker swap conditions. PPI greater than 0 indicate detection of the speaker swap prepulse. Circles indicate group mean ± standard error. Asterisks indicate the angle at which the population mean response is significantly different from 0 (* = p<0.05). (**f**) Prepulse inhibition (PPI) as a function of speaker swap angle of high-pass noise (<2 kHz) for Controls (top panel, blue) and CHL animals (bottom panel, red). (**g**) Prepulse Inhibition (PPI) with speaker swap angles combined using the broadband noise prepulse. The minimum audible angle (MAA) is defined as the the smallest angle where the PPI is significantly different from 0. Vertical dotted lines indicate the MAA for Control and CHL animals. For broadband noise, Control and CHL animals can both discriminate speaker swaps of ≥7.5°. (**h**) Prepulse Inhibition (PPI) as a function of combined speaker swap angles using the high-pass noise prepulse. For high-pass noise, Control animals can discriminate speaker swaps between 7.5° - 15°, whereas CHL animals discriminate between 30° - 45°.

Prior to testing with the speaker swap paradigm, baseline startle responses were characterized for all animals to determine whether Control and Early CHL animals displayed comparable startle responses. Figure 6d shows the mean startle response amplitude (± SEM) as a function of increasing sound intensity (dB) of the startle speaker in the presence of 70 dB background noise for controls (n=17) and animals raised with a CHL (n=25). An ANOVA revealed a significant difference between the two means (controls vs CHL groups) at one or more of the levels (one-way ANOVA with Bonferroni correction, F_2,627_ = 243.7, p<0.01), and a post-hoc student’s t-test found that the startle responses were significantly different at 80 dB (t(40)=-2.26, p=0.03) but not at 90 dB or above (90 dB: t(40)=-1.5, p=0.15; 100 dB: t(40)=-0.18, p=0.86; 110 dB: t(40)=-0.48, p=0.64; 120 dB: t(40)=-1.01, p=0.32). While there was a difference observed at 80 dB, there was no significant difference for sound intensities used for the speaker swap experiments (110-120 dB). Therefore, both groups exhibited comparable baseline startle responses to the eliciting stimulus prior to testing on the speaker swap paradigm.

The minimum audible angle in the frontal field was assessed by swapping the location of a continuous broadband or high-pass noise symmetrically across the midline between speakers separated by ±90°, 45°, 30°, 15°, and 7.5° (from the left-to-right and from the right-to-left) preceding the startle-eliciting stimulus (ES; 300 ms ISI). Control conditions, in which no swap occurred, were presented from each of the five starting speaker locations (i.e., noise presented from speaker 1, but did not swap to a second speaker prior to ES). Figure 6e, f shows the results of varying speaker swap angle across midline, where the PPI (population mean± SEM) for each angle condition is shown as a function of angle for broadband or high-pass noise speaker swaps for Control animals (top panels) and Early CHL animals (bottom panels). A PPI value of zero indicates that the startle response for the control condition (e.g., no speaker swap) is equal to the startle response with the prepulse present (e.g., speaker swap present), indicating that the swap in angle was *not detected.* Thus, PPI values that are significantly different than zero indicate that the animal *was* able to detect the speaker swap. To assess significance, a one-way ANOVA was used to determine angles that were significantly different from control, followed by Bonferroni-corrected two-tailed Student’s t-tests to identify specific angles that resulted in a significant change relative to control. The smallest angle that is significantly different from zero is defined as the minimum audible angle (MAA; cf. Mills 1958).

For the broadband noise condition (Figure 6e), which affords access to both low-frequency ITD and high-frequency ILD cues, an ANOVA revealed significant differences across angle for Controls (F_9,990_ = 6.03, p= 2.93×10^-8^). Post hoc analysis with Bonferroni correction revealed that PPI was significantly different from zero (i.e., compared to control no prepulse conditions) for left-to-right swaps in speakers separated by 7.5°, 15°, 30°, 45°, and 90° (all are: p<2.0735×10^-5^, df=99, t>4.47), and for right-to-left swaps separated by −15°, −30°, −45°, and −90° (all are: p<1.90×10^-9^, df =99, t>6.61). Figure 6g shows the left-to-right and right-to-left speaker swap angles collapsed (i.e., absolute value of speaker angle). The responses are significantly different at all angles tested (**7.5°**: p=0.004, df=199, t=2.92; **15°**: p=2.16×10^-14^, df=199, t=8.25; **30°**: p=3.94×10^-22^, df=199, t=10.95; **45°**: p=2.73×10^-39^, df=199, t=16.56; **90°**: p=6.14×10^-49^, df=199, t=19.8). Thus, for Controls, the MAA for broadband noise is less than 7.5°. For Early CHL animals, the ANOVA also revealed significant differences across angle (F_9,2490_ = 6.12, p= 1.5×10^-8^). Post hoc analysis with Bonferroni correction revealed that PPI was significantly different from zero for left-to-right speaker swaps of 7.5°, 15°, 30°, 45°, and 90° (all are: p<8.65×10^-4^, df=249, t>3.37), and for right-to-left swaps separated by −7.5°, −15°, −45°, and −90° (all are: p<9.31×10^-6^, df =249, t>4.52). Collapsing the left-to-right and right-to-left speaker swap angles, the responses are significantly different at all angles tested (**7.5°**: p=3.95×10^-11^, df=499, t=6.76; **15°**: p=4.89×10^-16^, df=499, t=8.39; **30°**: p=5.28×10^-4^, df=499, t=3.49; **45°**: p=4.14×10^-35^, df=499, t=13.38; **90°**: p=2.23×10^-78^, df=499, t=22.59). Thus, for early CHL animals, the MAA for broadband noise is also less than 7.5°, similar to the MAA for Controls. This suggests that for the broadband noise condition, Early CHL did not alter spatial acuity in the frontal field as compared to Controls.

For the high-pass noise condition (Figure 6f), an ANOVA revealed significant differences across angle for Controls (F_9,990_ = 5.86, p= 5.5× 10^-8^). Post hoc analysis with Bonferroni correction revealed that PPI was significantly different from zero (i.e., compared to control no prepulse conditions) for left-to-right swaps in speakers separated by 15°, 30°, 45°, and 90° (all are: p<2.66×10^-6^, df=99, t>4.98), and for right-to-left swaps separated by −30°, −45°, and −90° (all are: p<7.32×10^-9^, df =99, t>6.32). Figure 6h shows the left-to-right and right-to-left speaker swap angles collapsed (i.e., absolute value of speaker angle). The responses are significantly different at angles greater than 15° (**7.5°**: p=0.98, df=199, t=-0.03; **15°**: p=1.02×10^-5^, df=199, t=4.53; **30°**: p=3.05×10^-21^, df=199, t=10.65; **45°**: p=9.8×10^-49^, df=199, t=19.73; **90°**: p=1.65×10^-54^, df=199, t=21.76). Thus, for Controls, the MAA for high-pass noise is between 7.5° and 15°. For early CHL animals, an ANOVA revealed significant differences across angle (F_9,2670_ = 10.24, p= 1.17×10^-15^). Post hoc analysis with Bonferroni correction revealed that PPI was significantly different from zero for left-to-right speaker swaps of 30° and 90° (both are: p<1.98×10^-5^, df=267, t>4.34), and for right-to-left swaps separated by −45° and −90° (both are: p<9.31×10^-20^, df =267, t>9.89). Collapsing the left-to-right and right-to-left speaker swap angles (Figure 6h), the responses are significantly different at angles greater than 30° (**7.5°**: p=0.015, df=535, t=-2.44; **15°**: p=0.43, df=535, t=0.78; **30°**: p=6.82×10^-5^, df=535, t=4.01; **45°**: p=1.5×10^-12^, df=535, t=7.25; **90°**: p=4.48×10^-30^, df=535, t=12.13). Thus, for Early CHL animals, the MAA for high-pass noise is between 15° and 30°, which is two to four times larger as the MAA of 7.5°-15° for Controls.

### Early unilateral hearing loss alters neural coding of interaural level difference (ILD) cues in the auditory midbrain

Since interaural level difference (ILD) cues to sound location largely depend on high-frequencies, and Early CHL animals display poorer spatial acuity for sounds comprised of high-frequencies, we next asked whether the spatial deficits could be attributed to impaired ILD processing in auditory neurons. Indeed, Early CHL animals displayed alterations in the binaural auditory brainstem response (ABR), of which the derived binaural component is generated by the lateral superior olive (LSO; Benichoux et al., 2018), the site that first integrates binaural information and encodes ILD cues (Tollin 2003). Further, the PPI of the acoustic startle is mediated by the auditory midbrain (i.e., inferior colliculus; Koch 1999), which receives direct inputs from the LSO and contains neurons that encode ILD (Irvine and Gago, 1990). Therefore, we directly assessed ILD coding in the inferior colliculus (IC), from same animals that displayed BIC (Figure 4) and behavioral spatial hearing deficits (Figure 6). We collected single-unit extracellular recordings from IC neurons from littermate Control and Early CHL animals. For Early CHL, recordings were made in both the IC contralateral and ipsilateral to the previously occluded ear. For each isolated IC neuron, the characteristic frequency (CF), rate-level function at CF, and binaural sensitivity to ILD was assessed. Neurons were selected for further analysis if they met a set of criteria for ILD sensitivity (see Methods).

ILD sensitivity was determined by varying the intensity of the tone stimulus (the CF of the isolated unit) presented to both ears through the insert earpieces (Figure 7a). The sound level at the contralateral ear was held constant (i.e., excitatory ear), and the level to the ipsilateral ear (i.e., inhibitory ear) was then varied ±30 dB (5 dB steps). Figure 7b shows an example rate-ILD function for an ILD-sensitive IC neuron. All raw rate-ILD data were fit with a four-parameter sigmoidal logistic function (Supplemental Figure 3a, b) and were included for further analysis if specific criteria were met (see Methods). The parameters include the maximum and minimum firing rate (spikes/second, s/s) of the rate-ILD functions, the 50% inflection point of the sigmoid fit (i.e., half-maximum of fit), the ILD dynamic range (i.e., range of ILD producing 10-90% of maximal firing), and the slope of the rate-ILD functions.

**Figure 7.**
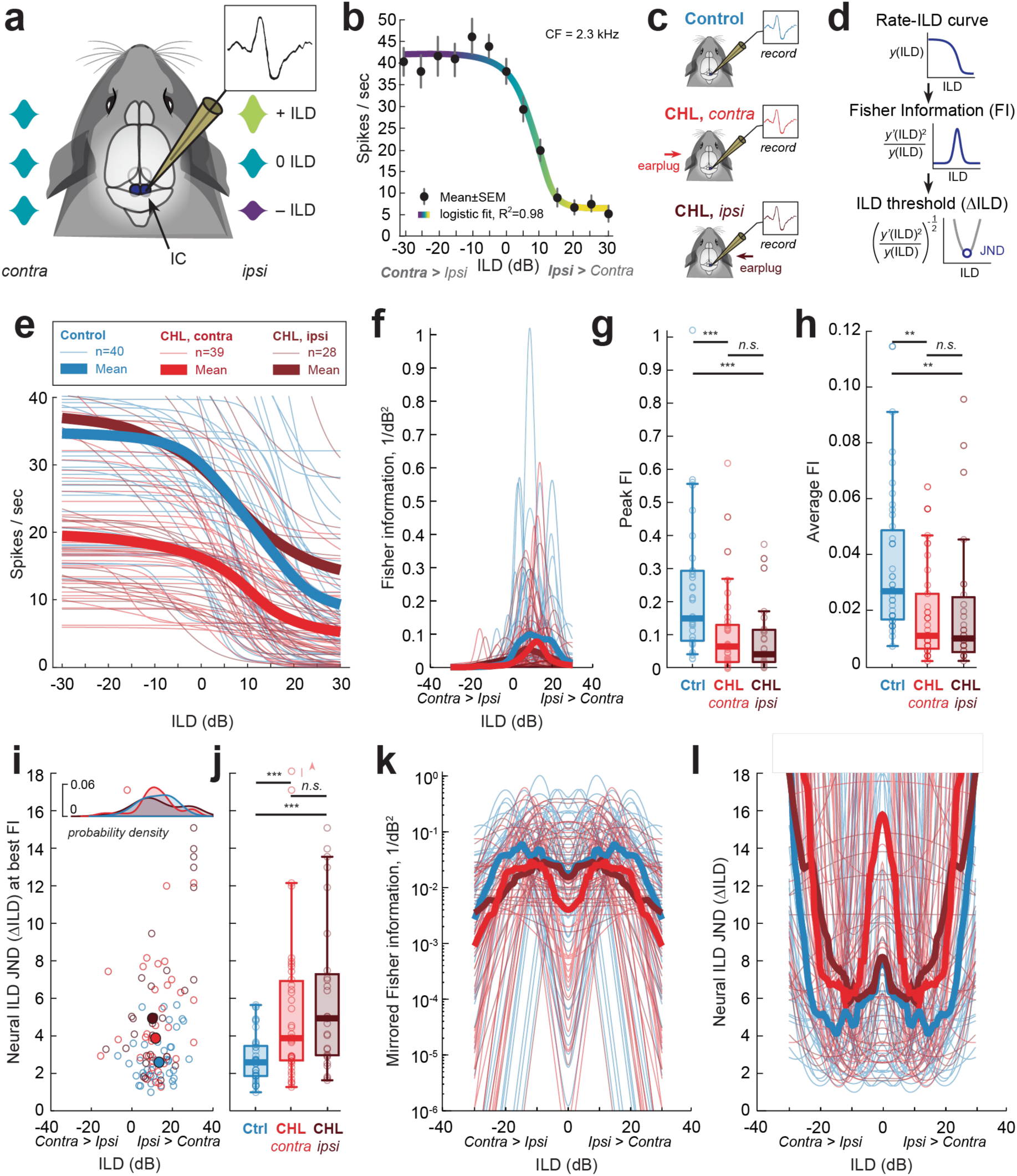
Unilateral developmental CHL degrades neural discrimination of interaural level difference (ILD) cues in auditory midbrain neurons. (**a**) Schematic of experimental approach. Single-unit extracellular recordings were collected from inferior collicular (IC) neurons and assessed for binaural sensitivity to interaural level difference (ILD) cues. The sound level of a tone (at the neuron’s characteristic frequency, 20 dB above threshold) is fixed at the ear contralateral to the recording site, and the sound level for the ipsilateral ear (i.e., providing the inhibitory drive) is varied (±30 dB, 5 dB steps). (**b**) Rate-ILD function for an example neuron that is responsive to ILD. Neurons are considered to be ILD-sensitive when firing rates are modulated by ≥ 50% with increasing sound level to the ipsilateral ear. (**c**) Overview of recording conditions for Control and CHL animals. Neurons recorded from the IC contralateral to the previous CHL are designated as “CHL, *contra*”. Neurons recorded from the IC ipsilateral to the previous CHL are designated as “CHL, *ipsi*”. (**d**) Rate-ILD curves from each neuron are used to compute Fisher information (FI), from which neural ILD thresholds (ΔILD) can be computed. The just-noticeable-difference (JND) for ILD is defined as the minimum of the ΔILD function. (**e**) Rate-ILD functions for single IC neurons from Control (blue; n=40), CHL contra (red; n=39), and CHL *ipsi* (dark red; n=28). Thick lines indicate group means, and thin lines indicate individual neurons. (**f**) Distribution of FI (1/dB)-½ as a function of ILD for Control, CHL contra, and CHL *ipsi* neurons. (**g**) The maximum FI for each distribution was computed for each unit. Control neurons displayed significantly higher peak FI values (median±SD: 0.15±0.2 1/dB2) than CHL *contra* (0.06±0.1 (1/dB2)), and CHL *ipsi* neurons (0.04±0.1 (1/dB2); p<0.001). (**h**) The average of the FI distribution for each unit. Control neurons displayed significantly higher average FI (0.03±0.2 (1/dB2)) than CHL *contra* (0.01±0.02 (1/dB2)), and CHL *ipsi* neurons (0.01±0.02 (1/dB2); p=0.002). (**i**) Distribution of neural ILD JNDs for Control (n=40), CHL *contra* (n=39), and CHL *ipsi* (n=28) single units. Open circles indicate individual neurons, and filled circles depict the median ILD location (x-axis) and corresponding ILD JND (y-axis) for each group (Ctrl: ILD location:13 dB ILD, median JND: 2.59 dB ILD; CHL *contra*: location: 11.3 dB ILD, median JND: 3.9 dB ILD; CHL *ipsi*: location: 9.8 dB ILD, median JND: 4.9 dB ILD). Top inset: probability density estimates for distribution of ILD location for each group (Ctrl, CHL *contra*, CHL *ipsi*). X-axis of distributions align with the x-axis for main plot in (**j**). Single IC neurons from Early CHL animals exhibit significantly elevated neural ILD JNDs compared to Controls (median±SD: Ctrl: 2.59±1.3 dB ILD; CHL *contra*: 3.9±4.2 dB ILD; CHL *ipsi*: 4.9±4 dB ILD; *** = p<0.001). (**k**) “Mirrored” population FI as a function of ILD. Each IC cell and corresponding rate-ILD function is assumed to have a mirrored equivalent in the opposite IC. The “left” and “right” IC cells are summed to compute the population FI that incorporates both hemispheres. (**l**) The neural ILD JND derived from the mirrored population FI in (k). The median (±SD) JND at midline (0 dB ILD) for Control cells is 7.8±164.4 dB ILD, for CHL *contra* cells is 15.8±396.1 dB ILD, and for CHL *ipsi* cells is 8.2±73.8 dB ILD. For all boxplots, the thick center line indicates the median, and the top and bottom of each box are the 75th and 25th percentiles, respectively.

The median (±SD) maximum firing rates of the ILD functions were 34.6 ± 20.9 s/s for Controls, 19.5 ± 9.8 s/s for CHL_contra_ neurons, and 27.6 ± 30.5 s/s for CHL_ipsi_ neurons (Supplemental Figure 3c). A Mann-Whitney U test revealed a significant difference between the maximum firing rate of Controls and CHL_contra_ neurons (z = 4.77, p < 0.0001) but not CHL_ipsi_ neurons (z = 0.66, p = 0.51). CHL_contra_ neurons were also significantly different from CHL_ipsi_ neurons (z =-2.75, p = 0.006). For the minimum firing rates of the ILD functions, the median (±SD) were 4.1 ± 13.6 s/s for Controls, 2.6 ± 4.73 s/s for CHL_contra_ neurons, and 4.6 ± 21 s/s for CHL_ipsi_ neurons (Supplemental Figure 3d). A Mann-Whitney U test found no significant differences between the groups (Controls vs CHL_contra_: z = 1.3, p = 0.19; Controls vs CHL_ipsi_: z = −0.36, p = 0.72; CHL_contra_ vs CHL_ipsi_: z = −1.44, p = 0.15).

For the 50% inflection point of the sigmoid fit, the median (±SD) were 10.4 ± 7.4 dB for Controls, 9.2 ± 9.8 dB for CHL_contra_ neurons, and 6.2 ± 9.0 dB for CHL_ipsi_ neurons (Supplemental Figure 3e). A Mann-Whitney U test found no significant differences between the groups (Controls vs CHL_contra_: z = 0.59, p = 0.56; Controls vs CHL_ipsi_: z = 1.96, p = 0.05; CHL_contra_ vs CHL_ipsi_: z = 1.61, p = 0.11). For the ILD dynamic range of the sigmoid fit, the median (±SD) were 15 ± 11.7 dB for Controls, 22.2 ± 13.8 dB for CHL_contra_ neurons, and 23.5 ± 16.5 dB for CHL_ipsi_ neurons (Supplemental Figure 3f). A Mann-Whitney U test revealed a significant difference between the ILD dynamic range of Controls and CHL_ipsi_ neurons (z = −2.99, p = 0.0028) but not CHL_contra_ neurons (z = −1.37, p = 0.17). There were no significant differences between CHL_contra_ and CHL_ipsi_ neurons (z = −1.65, p = 0.099). Finally, the slope of the rate-ILD functions were found to be significantly smaller in both CHL groups compared to Controls (Supplemental Figure 3g). The median slopes (±SD) were 3 ± 3.6 sp/s/dB for Controls, 0.6 ± 1.15 sp/s/dB for CHL_contra_ neurons, and 0.58 ± 3.4 sp/s/dB for CHL_ipsi_ neurons. A Mann-Whitney U revealed significant differences between Controls and CHL groups (Controls vs CHL_contra_: z = 5.2, p < 0.00001; Controls vs CHL_ipsi_: z = 3.28, p = 0.001) and no significant differences between CHL_contra_ and CHL_ipsi_ neurons (z = - 0.43, p = 0.67).

In summary, Early CHL animals exhibit the following differences in rate-ILD functions of midbrain neurons: (1) a reduction in maximum firing rate for both CHL_contra_ and CHL_ipsi_, (2) an increase in ILD dynamic range (dB) for CHL_ipsi_, and (3) shallower ILD slopes (sp/s/dB) for both CHL_contra_ and CHL_ipsi_.

### Early unilateral hearing loss degrades neural discrimination of interaural level difference (ILD) cues in the auditory midbrain

Other developmental plasticity studies of the auditory system have also reported changes to rate-ILD coding in the auditory midbrain (Silverman and Clopton 1977; Clopton and Silverman 1977; Popescu and Polly 2010). But, it is uncertain how these changes relate to a behavioral function. Here, using the startle-based behavioral paradigm, Early CHL animals displayed impairments with discriminating sound sources that rely on ILD cues. We employed the mathematical framework of Fisher Information (FI) to compute neural ILD discrimination values of IC neurons (Seung and Sompolinsky, 1993; Jones et al., 2015; Brown & Tollin, 2016; Benichoux et al., 2017; Brown et al., 2018), enabling us to compare neurophysiological and behavioral discrimination of sound sources.

Figure 7e shows rate-ILD functions for all IC neurons collected from littermate Controls (n=40) and CHL animals (contra: n=39; ipsi: n=28) that met the criteria for ILD sensitivity (see Methods). The distribution of FI is drawn from the rate-ILD functions (e.g., see Figure 7d). Figure 7f shows the distribution of FI across ILDs (±30 dB ILD) for ILD sensitive IC neurons. The maxima for the FI distribution for each unit was taken and compiled in a summary boxplot (Figure 7g). The median FI for CHL_contra_ neurons is 0.06 ± 0.1 (SD) 1/dB^2^, 0.04 ± 0.1 for CHL_ipsi_ neurons, and 0.15 ± 0.2 for Control neurons. We find both CHL groups exhibit a significant reduction in peak FI values compared to controls (Mann-Whitney U, controls vs CHL_contra_: z = 3.9, p = 0.0001; Controls vs CHL_ipsi_: z = 4.04, p < 0.00001). Peak FI between the two CHL groups was not significantly different (z = 0.74, p = 0.46).

Next, we quantified neural ILD sensitivity by computing neural ILD discrimination thresholds (i.e., the smallest change in ILD that a neuron can detect, given a specific criterion). FI can be directly related to the well-known discriminability index, *d’*, a signal detection metric used to quantify discrimination of sensory stimuli (Green and Swets, 1966; Seung and Sompolinsky, 1993; Tollin et al., 2008; Jones et al., 2015b; Brown and Tollin, 2016b; Benichoux et al., 2017). Here, we use FI to evaluate the change in ILD necessary to achieve 1 standard separation in spiking distributions (i.e., the neural just-noticeable difference, JND, for a *d’* = 1). In this formula, higher FI values correspond to lower neural ILD thresholds (i.e., better ILD discrimination). Figure 7i shows the ILD JNDs across ILDs (±30 dB ILD) for Control and Early CHL neurons. The average neural ΔILD for control neurons is 2.6 ± 1.3 dB ILD (median ± SD), for CHL_contra_ neurons is 3.9 ± 4.2 dB ILD and for CHL_ipsi_ neurons is 4.9 ± 4 (Figure 7j). We find both Early CHL groups exhibit significantly elevated ILD JNDs (∼1-2 dB ILD worse) as compared to controls (Controls vs CHL_contra_: z = −3.9, p = 0.0001; Controls vs CHL_ipsi_: z = −4.04, p < 0.00001). Early CHL groups were not significantly different from each other (z = −0.74, p=0.46).

Assuming that ILD discrimination depends on neural populations from both left and right IC nuclei, we next opted to pool the data to reflect FI and JND that incorporates both hemispheres (Figure 7k,l). Each IC cell and corresponding rate-ILD function is assumed to have a mirrored equivalent in the opposite IC. The “left” and “right” cells are summed to compute the population FI that includes both hemispheres (“Mirrored FI”; Figure 7k). Both CHL_contra_ and CHL_ipsi_ neurons display lower FI at non-zero ILDs compared to Control neurons. The median FI across all ILDs (±30 dB ILD) were lower in CHL neurons (Control: 0.03; CHL_contra_: 0.011; CHL_ipsi_: 0.012). At midline (0 dB ILD), CHL_contra_ neurons have lower FI than CHL_ipsi_ and Controls (Control: 0.018; CHL_contra_: 0.004; CHL_ipsi_: 0.015). The neural ILD JND is then recomputed based on the Mirrored FI to generate the population JND that represents neurons from both IC nuclei (Figure 7l). Both CHL neurons displayed poorer ILD JNDs for non-zero ILDs. The median JND across all ILDs (±30 dB ILD) were elevated in CHL neurons (Control: 5.9; CHL_contra_: 9.3; CHL_ipsi_: 9.1). At midline, CHL_contra_ neurons had poorer JNDs at midline compared to CHL_ipsi_ and Control neurons (median±SD: Control: 7.8±164.4 dB ILD; CHL_contra_: 15.8±396.1 dB ILD; CHL_ipsi_: 8.2±73.8 dB ILD).

In summary, we find that after a transient developmental CHL in one ear, ILD-sensitive neurons in the auditory midbrain display a reduction in sensitivity to ILD compared to littermate Controls. Further, these animals displayed altered binaural processing in the auditory brainstem along with deficits in spatial discrimination of sounds that largely contributes to ILD cues (i.e., high-frequencies).

### Early unilateral CHL does not alter monaural response properties in the auditory midbrain

One potential explanation for altered ILD processing by IC neurons is that monaural response properties may be compromised following Early CHL. Therefore, we examined monaural frequency tuning curves, rate level functions, and first spike latencies (see Methods) for all ILD-sensitive neurons.

Supplemental Figure 4a shows an example “response area curve” (a 3D surface plot of frequency × intensity × firing rate) from which the neuron’s characteristic frequency (CF) can be determined (e.g, see asterisk). Supplemental Figure 4b shows the neuron threshold as a function of unit CF for controls (n=40) and Early CHL animals (CHL_contra_, n=39; CHL_ipsi,_ n=28). Histograms for CF and threshold were plotted on the x- and y-axes, respectively, to aid in visualization of data distribution. The mean neuron threshold (± SD) was 45.88 ± 13.4 dB for controls, 52.4 ± 11.5 dB for Early CHL_contra_, and 49.82 ± 12.9 dB for Early CHL_ipsi_. A one-way ANOVA found no significant differences between the threshold means (F_2,104_ = 2.71, p = 0.071). The CFs between the groups displayed similar ranges in frequency (Controls: 0.86-25.73 kHz; CHL_contra_: 0.649-15.91 kHz; CHL_ipsi_: 0.559-26.86 kHz), although neurons ipsilateral to the CHL had a higher proportion of high CF units (>8 kHz). The range of CFs observed are comparable to the guinea pig behavioral audiogram reported in Heffner et al. (1971).

Next, we examined the rate level functions (RLFs) from all neurons. Once the CF is determined for each unit (from the response area curve), the intensity at the contralateral ear (the excitatory ear) is varied to determine unit threshold and first spike latency to CF tone. Supplemental Figure 5a shows an example RLF from an IC neuron (left panel). The right panel shows the associated spike rasters for each trial, at each sound intensity presented (30-90 dB at the contralateral ear). The dotted line indicates the onset of the spike responses (i.e., first spike latency). Supplemental Figure 5b shows the first spike latency for Control and Early CHL neurons as a function of CF. The mean latency (± SD) for controls is 17.28 ± 5.38 ms, for early CHL_contra_ is 17.4 ± 3.54 ms, and for early CHL_ipsi_ is 17.12 ± 6.92 ms. A one-way ANOVA revealed no significant differences between the first spike latency between Control and Early CHL neurons (F_2,104_ = 0.03, p = 0.98).

In summary, monaural response properties were not altered following developmental CHL as compared to Controls. Therefore, we attribute the altered ILD coding and poorer neural ILD JNDs following Early CHL to alterations to binaural, and not monaural, processing in IC neurons.

## Discussion

Experience-driven developmental plasticity of binaural processing has been widely examined at the level of the auditory thalamus (Miller and Knudsen, 2003) and cortex (Brugge et al., 1985; Popescu and Polley, 2010; Keating et al., 2013a, 2015; Polley et al., 2013). Fewer studies, however, have examined whether developmental auditory experience can also induce plasticity at earlier ascending nuclei, including the auditory brainstem, the initial site of synaptic convergence between input from both ears (Kandler et al., 2009), or the auditory midbrain (inferior colliculus, IC) (Silverman and Clopton 1977; Clopton and Silverman 1977; Popescu and Polly 2010; Thornton et al 2021; see Kumpik and King 2019 for review). Here, we report a prolonged maturation of the binaural auditory brainstem in guinea pigs, suggesting that binaural plasticity may be heightened, and therefore vulnerable to experience, during this time. Indeed, we find that developmental HL alters a brainstem readout of binaural function which is not observed when the HL is induced in adulthood. Startle-based behavioral measures reveal poorer spatial resolution of sound sources, but only for high-frequency sound stimuli. Finally, single-unit recordings of auditory midbrain neurons reveal significantly poorer neural acuity to a sound location cue that largely depends on high-frequency sounds.

### Altered binaural processing in the auditory brainstem following developmental hearing loss

Following developmental CHL, we find alterations of the binaural interaction component (BIC) of the auditory brainstem response (ABR), including longer latencies and broader “tuning” to ITDs. Our findings are consistent with reports in both human and animal subjects. Laska et al. (1992) found similar increases in BIC DN1 peak latency after rearing guinea pigs with a reversible CHL. They report latency differences of ∼0.1-0.4 ms which is comparable to that found in our study (∼0.13 ms). This study and others report longer latencies of later ABR waves that correspond to binaural auditory structures in children with a history of CHL (Folsom et al., 1983; Gunnarson and Finitzo, 1991; Hurley and Hurley, 1995).

An increase in BIC latency suggests that a developmental CHL can lead to temporal alterations in the binaural nuclei of the auditory brainstem. There are several potential contributions that may increase the latency of ABR waves, including increased conduction time between auditory nuclei in the ascending pathway or reduced temporal coding (i.e., synchronicity of neurons to stimulus). Myelination of axons, which contributes to conduction velocity, is an activity-dependent process (Fields, 2015). In fact, acoustic trauma can result in demyelination of the auditory nerve resulting in slower conduction velocities (Ito et al., 2004). Given that precise timing of axonal inputs is critical for brainstem processing of acoustical cues (Brown and Tollin, 2016b; Beiderbeck et al., 2018; Franken et al., 2018), asymmetrical CHL in the developing brainstem may lead to axon conduction imbalances between synaptic inputs from the left and right ear.

The BIC is strongly modulated by acoustical cues to sound location, including ITDs. In the present study, we find developmental CHL leads to decreases in the BIC amplitude at small ITDs, reduced modulation of BIC amplitude to ITD, and broader BIC vs ITD tuning functions. The decrease in BIC amplitude suggests that there is less inhibition when comparing the binaural-evoked ABR with the sum of the monaural ABRs (the BIC is a negative peak, so the greater the inhibitory drive, the smaller the binaurally-evoked ABR, the larger the BIC peak). This suggests a CHL-induced disruption of the developing inhibitory binaural circuits. A reduction in BIC amplitude could also suggest less synchronous activity by neurons in the circuit. Broader BIC vs ITD curve widths could indicate that neurons in the E/I circuit are not as precisely tuned, requiring larger changes in ITD to synchronously modulate the BIC amplitude. Animals with asymmetrical acoustic experience during development could result in imbalances in excitation and inhibition. In fact, Clarkson et al. (2016) found that a monaural developmental CHL in rats led to an upregulation of AMPA glutamate receptor subunits (GluA3) on bushy cells of the cochlear nucleus, and AMPA receptors mediate fast synaptic transmission in the auditory pathway. In the cochlear nucleus, inhibition is mediated by the activation of glycine receptors (Hirsch and Oertel, 1988), which are reported to be downregulated in the cochlear nucleus following CHL (Whiting et al., 2009). Up- or down-regulation of excitatory and inhibitory neurotransmitter receptors reflects compensatory homeostatic mechanisms in neuronal circuits following external disruptions (Wierenga et al., 2005), and can alter the efficiency of synaptic transmission (Manilow and Malenka, 2002).

The developing auditory brainstem, including the LSO, is highly influenced by activity-driven modifications after the onset of hearing (Sanes and Rubel, 1988; Sanes and Wooten, 1987). Indeed, we find that the BIC responses in newborn guinea pigs are poorly tuned to ITD and undergo a period of prolonged refinement. Further, the rate at which the BIC tuning improves aligns with the rate at which the head diameter grows (Anbuhl et al., 2017) suggesting that the time course of BIC maturation parallels the time course of head growth and the resulting changes in magnitude of the acoustical cues to sound location. Given that auditory neurons in the brainstem must maintain rapid transmission and temporal fidelity (Trussell, 1999), diminished acoustic experience in one ear could disrupt this maturation process leading to changes in the E/I balance in the brainstem.

### Spatial hearing deficits following developmental hearing loss

Human auditory perceptual skills mature over a long time, such as frequency resolution which matures by 6 months of age (Hall and Grose, 1991), to frequency discrimination (i.e, discriminating a difference between two tones presented sequentially) maturing as late as ∼10 years of age for low-frequency sounds (Moore et al., 2011). Intensity discrimination (i.e., distinguishing whether one sound is louder than the other) does not mature until after 10 years of age (Maxon and Hochberg, 1982). Infants can detect differences in intensity between two sounds of about 6 dB, and this decreases to about 2 dB by 4 years of age (Sinnott and Aslin, 1985). By comparison, adults can discriminate intensity differences of high frequency tones as little as 0.5 dB (Mills, 1960). Children that experience CHL during these developmental ages often have long lasting binaural hearing deficits that can persist for years beyond resolution of the CHL (Hogan and Moore, 2003; Koiek et al., 2022; Ludwig et al., 2019; Mishra & Moore 2023; Moore et al., 1991; Nittrouer & Lewenstein 2024; Pillsbury et al., 1991).

Animal studies also report persistent perceptual deficits following developmental hearing loss. In a classic series of experiments, barn owls were subjected to unilateral CHL (i.e., earplug in one ear) either during development or as adults. When tested in an auditory orienting (head turn) paradigm after plug removal, both groups initially mislocalized the source in the direction of the previously occluded ear. However, the adult-CHL animals adapted to the corrected binaural inputs, while the developmental-CHL animals continued to mislocalize the source (Knudsen et al., 1984a, 1984b). Similar observations have been reported for localization and tone-in-noise detection tasks in ferrets (Keating et al. 2013; King et al. 2001; Moore et al. 1999). Together, these studies suggest the existence of a binaural auditory “sensitive period” (Knudsen et al., 1984a, 1984b), beyond which amelioration of peripheral inputs may be ineffective in correcting aberrant central mapping.

Here, we find that Early CHL animals required larger changes in the location of high-pass noise to detect a change in spatial location. The broadband and high-pass stimuli conditions correspond to the relevant acoustical cues available to the guinea pig (broadband: ITDs and ILDs; high-pass: ILDs and spectral cues). Early CHL animals required larger minimum audible angles (MAA) to discriminate changes in source location compared to control animals for high-pass noise only, suggestive of an impairment in ILD acuity. To estimate the deficit in behavioral ILD sensitivity, acoustic transfer function measurements reported for guinea pigs were utilized (Greene et al., 2014). Here, the average ILD vs azimuth slope (dB/degree) was computed for sources ±30° about the midline based on a population of adult animals. For frequencies greater than 4 kHz (the frequency cutoff for our high-pass noise stimuli), the average ILD slope is ∼0.2 dB/°. Using the high-pass noise behavioral discrimination thresholds (MAAs), the MAA can be multiplied by the average ILD slope to get an estimation of behavioral ILD sensitivity. The MAA for controls was between 7.5-15°, so the estimated ILD sensitivity is 7.5-15° × 0.2 dB/° = 1.5-3 dB ILD. The MAA for early CHL animals was between 15-30°, so the estimated ILD sensitivity is 15-30° × 0.2 dB/° = 3-6 dB ILD. Thus, we estimate that CHL we induced in developing guinea pigs impaired ILD sensitivity by ∼1.5-3 dB. Animals also displayed behavioral deficits, along with alterations in the latency and amplitude of the BIC DN1 peak of the ABR. This is consistent with the human literature, which finds that changes in the BIC with ILDs or ITDs is predictive of psychophysical performance in lateralization, discrimination, and binaural masking level difference tasks in both typical and hard-of-hearing individuals (Furst et al., 1990; Hall and Grose, 1993a,b; Riedel and Kollmeir, 2002a).

### ILD-sensitive midbrain neurons from early CHL carry less information regarding ILD, leading to impairments in ILD discrimination

The BIC and the behavioral results indicate that there are alterations in underlying binaural processing of ILD cues at the earliest levels of the auditory system. The initial processing of ILD cues occurs in the lateral superior olive (LSO; Goldberg and Brown, 1968, 1969; Tollin, 2003). The LSO projects to the central nucleus of the inferior colliculus (IC), a crucial center for spatial hearing. Lesions of the IC result in severe deficits in localization tasks, but not necessarily in other basic auditory tasks (e.g., Jenkins and Masterton, 1982; Kelly and Kavanagh, 1994). The effect of altered inputs to the IC may also be manifested in clinical studies of children with histories of early CHL, who often exhibit altered BICs of the ABR that are predictive of impaired spatial hearing (Gunnarson and Finitzo, 1991). The latencies of the BIC indicate a brainstem-level deficit in binaural processing occurring by the level of the IC (Møller, 2007; Benichoux et al., 2018; Tolnai and Klump 2020). Given the difficulty of directly recording from ILD-sensitive cells in the LSO (i.e., the neural generator of the BIC (see Owrutsky et al., 2021 for review)), we opted to record from neurons in the inferior colliculus which receives direct LSO inputs (Tsai et al., 2010). Furthermore, the IC is required for the PPI of the acoustic startle (Koch, 1999), implying that any disruption in encoding of the cues to location by neurons in the IC would also disrupt the PPI of the acoustic startle to changes in sound location.

Unilateral CHL during development can lead to extensive changes to auditory brainstem anatomy (Webster, 1983; Moore et al., 1989) and physiology (Silverman and Clopton, 1977; Moore and Irvine, 1981; Knudsen, 1999; Popescu and Polley, 2010; Thornton et al., 2021). The neurophysiological changes that are observed in many of these studies suggest an imbalance of converging ipsilateral and contralateral inputs to the IC. For example, experiments conducted by Silverman and Clopton (1977) showed that rearing rats with a unilateral CHL resulted in substantially reduced effective inhibitory input to the IC ipsilateral to the occlusion, and a considerable increase in effective inhibitory input to neurons in the contralateral IC. Similar results have been reported in the IC and cortex of the rat (Popescu and Polley, 2010) and mouse (Polley et al., 2013). Opposite results (increased ipsilateral inhibition, decreased contralateral inhibition) have been reported in the cat (Moore and Irvine, 1981), barn owl (Mogdans and Knudsen, 1992) and chinchilla (Thornton et al., 2021), perhaps indicative of species differences. All studies, however, demonstrated that CHL led to disruptions in inhibition and impaired neural ILD sensitivity that persist beyond resolution of the CHL.

Here, we observed differences in ILD sensitivity between IC neurons from normal and Early CHL animals. We found a reduction in maximum firing rate between Controls and both Early CHL groups (ipsi, contra to CHL), suggesting that the excitatory drive to the ear both contralateral and ipsilateral to the previous CHL is lessened. While there were no significant differences in the minimum firing rates, a proportion of neurons ipsilateral to the CHL (which largely provides the inhibitory drive to the IC) appear to have higher minimum firing rates, suggesting a lessened inhibitory input. For the 50% inflection point (or the half-maximum value) there were no significant differences between groups, although there appears to be a “shift” in the inflection point towards negative ILD values indicating that a proportion of neurons is now encoding a different range of ILDs. For the ILD dynamic range, there is a significant increase in the dynamic range for neurons ipsilateral to CHL compared to controls, and a similar trend for neurons contralateral to CHL. There appears to be less effective inhibitory influences from the ear ipsilateral to the CHL resulting in neurons encoding larger ranges in ILD compared to control neurons, which are encoding a narrower range of ILDs. For the ILD slope, the smaller values for CHL neurons indicates that the slope of the function is much shallower compared to control neurons. All else being equal, a shallowing of the rate slope indicates that the neurons carry less information about ILD (Thornton et al., 2021). Overall, the results are suggestive of an alteration in the excitatory/inhibitory input to the IC in early CHL animals, leading to possible reductions in ILD sensitivity.

We opted to use a more quantitative approach to assess the impacts of CHL on neural coding: the mathematical framework of Fisher information (FI). FI quantifies how well a neuron can discriminate two stimulus values (e.g. two ILDs), thus measuring stimulus precision. FI is a useful metric that allows for the quantification of neural ILD discrimination− which, importantly, is a more direct comparison between neurophysiological responses and behavioral performance as they both measure discrimination of sound source locations. There were reductions in FI between Early CHL and control neurons with a ∼53% reduction for neurons contralateral to the CHL and a ∼73% reduction for neurons ipsilateral to the CHL. Using FI, it was determined how well neurons could discriminate changes in ILD (i.e., neural ILD JND): early CHL neurons could discriminate changes of ∼4-5 dB ILD which is twice that of control neurons (ILD JND: ∼2.6 dB). Comparing the neural ILD JNDs, early CHL impaired ILD sensitivity of ICC neurons by ∼1.4-2.4 dB. The ILD discrimination thresholds obtained from the behavior and physiology both show similar impacts of early CHL. For Control animals, the behavioral ILD threshold was estimated to be 1.5-3 dB ILD, while the average ILD JND for single neurons was ∼2.6 dB ILD. The JNDs from Early CHL animals were also consistent, with a behavioral ILD JND estimated between 3-6 dB ILD and a neural ILD JND of ∼4-5 dB ILD. Thus, both behavioral and physiological measures of ILD JNDs show at least a *two-fold increase* (i.e., twice as impaired) following a developmental CHL compared to Controls.

### Conclusions

While animal studies have examined the anatomical, physiological, and behavioral implications of developmental CHL on binaural processing of the auditory thalamus and cortex, it was still unclear whether unilateral deprivation can also induce plasticity in the auditory brainstem. Here, we report a prolonged maturation of the binaural auditory brainstem in the guinea pig by tracking auditory evoked potentials across development. Using this age range, we induced a reversible unilateral HL and asked whether behavioral and neural maturation were disrupted. We found that developmental HL altered a brainstem readout of binaural function which was not observed when the HL was induced in adulthood. Startle-based behavioral measures revealed poorer spatial resolution of sound sources, but only for high-frequency sound stimuli. Finally, single-unit recordings of auditory midbrain neurons revealed significantly poorer neural acuity to a sound location cue that largely depends on high-frequency sounds. Taken together, these findings show that unilateral deprivation can disrupt developing auditory circuits that integrate binaural information and may give rise to the lingering spatial hearing deficits observed in human and animal studies.

## Methods

### Experimental subjects

A total of 85 guinea pigs (*Cavia porcellus*) (44 females, 41 males) were used in the study. Animals were used either for the Developmental ABR/BIC experiments or the conductive hearing loss (CHL) experiments. All pups were from our in-house breeding colony. Animals were housed on a 12-hour light / 12-hour dark cycle with full access to food and water. All surgical and experimental procedures complied with the guidelines of the University of Colorado Anschutz Medical Campus Animal Care and Use Committees and the National Institutes of Health.

### Developmental ABR/BIC experiment

#### Animal preparation

A total of 18 pigmented guinea pigs from an in-house breeding colony were used for ABR assessments across development (9 females, 9 males). Seven animals were used to track ABRs from birth (P1) through adult ages (>P56). The remaining eleven animals had 1-2 ABR measurements at different developmental ages. Prior to testing, animals were anesthetized intraperitoneally using a mixture of ketamine hydrochloride (<P14: 65-70 mg/kg; >P14: 80 mg/kg) and xylazine hydrochloride (<P14: 7 mg/kg; >P14: 8 mg/kg). The core body temperature was maintained by an electronically controlled heating blanket. Heart rate, respiration rate, and blood-oxygen levels (SpO_2_) were monitored throughout each experiment.

#### Experimental setup

The experimental details are briefly described here as a more in-depth description can be found in Ferber et al. (2016). Stimuli were presented through custom stainless-steel insert earpieces using TDT System 3 (Tucker Davis Technologies, Inc., Alachua, FL) CF1 speakers powered by a TDT SA1 amplifier. Sound level and phase were calibrated for each session via Etymotic Research (Elk Grove Village, IL) ER-7C microphones, with probe tubes positioned ∼2 mm into the ear canal. The calibration was applied using a 129-tap minimum phase filter, which ensures delivery of a temporally discrete click (see Beutelmann et al. 2015 for more details). The microphones were kept in place during the experiment, and their output was recorded simultaneously with the ABR signal to ensure signal fidelity throughout the duration of each experiment. The absolute sound pressure level was referenced to a 1-kHz tone. Recordings were made with platinum subdermal needle electrodes (F-E2-12 electrodes; Grass Technologies, West Warwick, RI) at the apex (active) and nape of the neck (reference) with a hind leg ground (Figure 2b). Before acquisition, sound-evoked ABR signals were preamplified (10,000x) and band-pass filtered (cutoffs 100 Hz and 3 kHz) by a WPI ISO-80 amplifier. An automated artifact rejection threshold, typically ∼15 µV, was set before each recording session.

#### Recording parameters and data acquisition

Click-evoked monaural ABR thresholds in each animal were assessed using clicks at increasing intensity (generally 20-90 dB in 5dB steps; click rate: 33/s). Five hundred artifact-free repetitions were presented for each condition in randomized order. Monaural- and binaural-presented clicks were then varied across an ITD range spanning ±2000 μs (±2000, 1000, 750, 500, 375, 250, 125, 0 μs; all conditions randomized), bandpass-filtered (100Hz-3kHz), and averaged. The range of ITDs presented here encompasses and exceeds the physiological range of ITD in guinea pig, which is ∼±320 μs (Greene et al., 2014). One thousand artifact-free repetitions were recorded for each condition. Presentation of stimuli and acquisition of evoked potentials were facilitated by a Hamerfall Multiface II (RME) sound card controlled by PC running custom MATLAB software in a Linux-based environment (OpenSUSE 13.1).

#### BIC computation, normalization, and Gaussian fitting

Binaural and summed ABR waves were identified as local maxima and BIC waves as local minima by a custom MATLAB procedure. The BIC waveform was calculated for each ITD (BIC = ABR_Binaural_ − [ABR_LeftMonaural_+ ABR_RightMonaural_]) with the left ear and right ear ABR traces time-shifted before summation according to the timing of clicks corresponding to the ITD (see Figure 2b-d). Normalization of BIC amplitude was applied to account for across-animal signal-to-noise variability in individual recordings (Ferber et al., 2016b). Here, the BIC amplitudes were normalized to the average root mean square (RMS) of the two monaural ABRs. A four-parameter Gaussian curve was fit to the BIC peak amplitude versus ITD functions: BIC_model_ (ITD) = B + A*exp(−0.5*((ITD-C)/ σ)^2^) where C is the ITD of maximal BIC (i.e., curve center), σ is the curve width of the Gaussian function, A is the modulation of the curve (e.g., amplitude), and B is the baseline BIC amplitude value (Figure 2g).

A Spearman’s rho correlation analysis was used to assess whether there were significant developmental changes in the data (e.g., in amplitude, latency, Gaussian fit parameters). The Spearman’s correlation coefficient, *r_s,_* measures the strength and direction of association between two ranked variables, such as between the parameter of interest (e.g., BIC amplitude) and age. When a significant correlation is found, an exponential decay function is fit to the data. The methods used here are derived from the appendix of Eggermont & Moore (2012) which describe the use of exponential functions to model maturational changes.

#### Excitatory-inhibitory model of the BIC

Ungan et al. (1997) proposed a computational model to explain the dependency of the BIC latency and amplitude with ITD by modeling the binaural interaction found in cells of the LSO, with contralateral inhibitory and ipsilateral excitatory inputs (IE cells). The Ungan model has 4 parameters, including the difference between the mean arrival times of the excitatory and inhibitory input to the binaural LSO cell, the standard deviations of the excitatory and inhibitory arrival times, and the duration of inhibition of the LSO cell. Later, Riedel and Kollmeier (2006) adjusted the model to include an amplitude scaling factor (i.e., the four parameters are optimized by means of a chi-squared fit of the average BIC amplitude and latency data over subjects). This model was further refined in Benichoux et al. (2018), which is the model presented below.

Here, we sought to simplify this model by reducing the number of parameters from 4 to 2 as we are not interested in the latency of DN1, but rather how the amplitude changes with ITD. Consider that spikes arrive at different times for the excitatory and inhibitory pathways:

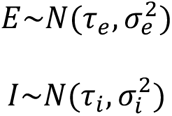

The binaural difference (i.e., the BIC) is assumed to measure the number of excitatory spikes that are cancelled by inhibition. This is obtained when excitation occurs after inhibition, within a given duration *w* (the inhibition *window*). Therefore, the event *R* occurring when a spike is cancelled by inhibition is defined as a function of the arrival times *I* and *E*, by:

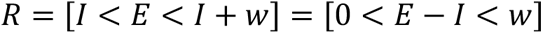

Where [x] is 1 if *x* is true and zero otherwise. Because the two random variables *E* and *I* are independent, normally distributed random variables, we know that the distribution of *U = E −I* is also normal:

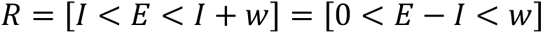

Since 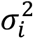 and 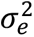 play an interchangeable role, we let 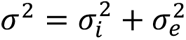, and for similar reasons, only the difference in mean arrival times plays a role, so we let 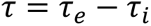 In turn, the probability that R is one is described with:

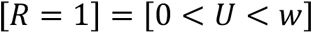

With 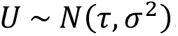. This probability is obtained with the cumulative function of a normal random variable. The probability that a spike gets canceled by inhibition is thus given by:

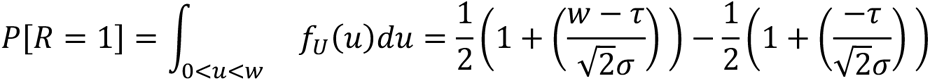

Where *f_u_*(*u*) is the probability density function of *U*. We can then identify the frequency of *R* = 1 to the magnitude of the BIC. Indeed, consider that the response of the population of LSO neurons is the summed response of *N* neurons: the mean rate 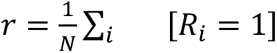. If we assume that all *R_i_* are independent and identically distributed, the response of the population is given by the frequency of *R* = 1, as given by the above equation. In order to obtain a model of the amplitude of the BIC, we have to consider LSO populations on both sides. We note that, on the left LSO, *τ_e_* = −*ITD*/2 and *τ_i_* = *ITD*/2 while on the right LSO, *τ_e_* = *ITD*/2 and *τ_i_* = −*ITD*/2. We are left to consider two normal random variables:

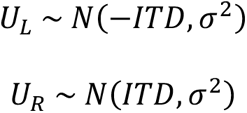

Whose responses follow from the previous equations:

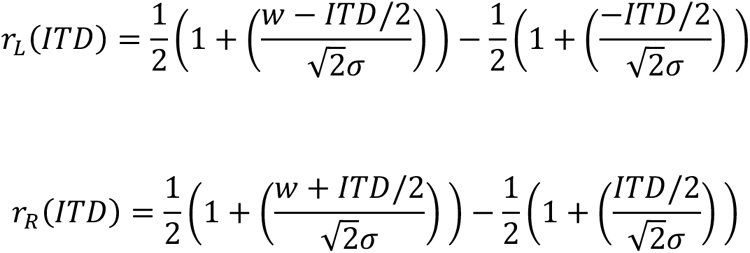

And, finally:

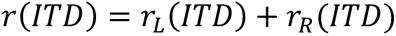

This simple expression provides us with a model of the BIC that depends on two parameters:

1. *σ* representing the precision of arrival times of excitatory and inhibitory inputs to LSO cells
2. *w* representing the temporal duration of the inhibition

The model predicts that the BIC should be roughly bell-shaped with a peak at 0 ITD. The width of the bell and its maximum depend on the relative values of ITDs, *σ*, and *w*. This model can then fit the BIC amplitude versus the ITD data. To do this, avoiding superfluous free parameters, the BIC amplitudes must be normalized between 0 and 1using the following expression:

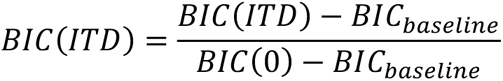

Where *BIC*(0) is the BIC at 0 ITD, and we obtain the baseline BIC using the BIC at very large ITDs (higher than 2 ms):

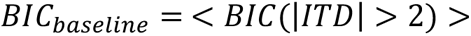

A least-squared fit can then be applied to the BIC amplitude values.

### Conductive hearing loss (CHL) experiments

#### Animal preparation

A total of 58 pigmented guinea pigs were used for CHL experiments. For developmental CHL, newborn animals from our breeding colony were divided into two groups shortly after birth (P0-P1): animals raised with normal hearing (controls, n=25; 11 males, 14 females) and animals raised with a unilateral CHL (designated as “Early CHL” throughout the paper; n=29; 11 males, 18 females). An additional 9 normal-hearing adults were used for physiology (8 males, 1 female) and were included in the Control physiology dataset. Animals were raised until adulthood, defined here as when the head and pinnae and hence the binaural acoustic cues to location reach adult dimensions, which occurs at postnatal day 56 (P56; i.e., 8 weeks after birth; Anbuhl et al 2017), after which the earplug, if present, was removed. The following were then assessed: 1) auditory brainstem responses (ABRs), 2) spatial discrimination via a startle-based approach, and 3) physiological recordings in the inferior colliculus. All testing was completed within 3-4 days of plug removal; see Figure 3c for a timeline of experiments.

A smaller group of adult guinea pigs (>P56; n=4) from our in-house colony were used to evaluate impacts of unilateral CHL in adulthood. For these experiments, baseline ABR assessments were completed (designated as “Pre-CHL”; 3 assessments per animal). after which a unilateral CHL of the same duration as that employed for the “Early CHL” group (8 weeks) was induced via earplugging. Following earplug removal, ABRs were collected for the post-CHL assessments (designated as “Post-CHL”; 2 sessions per animal; see Figure 5a for a timeline of experiments).

Prior to physiological testing (ABRs for Early CHL and adult CHL groups; single unit recordings for Early CHL groups), animals were anesthetized with an intraperitoneal injection of ketamine hydrochloride (80 mg/kg) and xylazine hydrochloride (8 mg/kg). Subsequent maintenance doses were administered when needed (60-70 mg/kg ketamine every hour, 4 mg/kg xylazine every other hour). The core body temperature was maintained by an electronically controlled heating blanket. Heart rate, respiration rate, and blood-oxygen levels (SpO_2_) were monitored throughout each experiment.

#### Earplug method of inducing CHL

Custom silicone earplugs (Microsonic Ready-press ear mold impression system; Westone Laboratories Inc, Colorado Springs, CO) were used to induce a transient, reversible conductive hearing loss (CHL). Prior to insertion of silicone material, the ear canal was cleaned with Epi-Otic Advanced Ear Cleaner to prevent infection. Plugs were placed in one ear within one day of birth (P0-P1) or in adulthood (>P56) and sealed with 3M™ Vetbond™ Tissue Adhesive. Care was taken to ensure that the silicone material did not go near the tympanic membrane, which can cause irritation and discomfort (see Figure 3a for approximate depth of plug within the ear canal). Plugs remained in place for 8 weeks, until developing animals reached adulthood (P56). We defined adulthood as the age at which the guinea pig head and external ear dimensions, which give rise to the binaural cues to sound location (ILD, ITDs), reach adult ranges (Anbuhl et al., 2017). Adult-onset CHL animals had earplugs in place for the same duration (8 weeks). Earplugs were checked 1-2x daily and replaced as needed.

Earplug attenuation was determined by placing a microphone (Type 4182, Bruel and Kjaer) deep within the ear canal of cadavers (similar to the methods described in Greene et al., 2014). A small puncture was made behind the pinnae in the mastoid region and the probe tip was inserted through to the ear canal; this was done so that the microphone would not be obstructing the delivery of the sound stimuli or the placement of the plugs). A free-field speaker presented a sweep in tone frequencies, from 100 Hz to 30 kHz (100 Hz steps) and the resultant dB SPL measured by the microphone was obtained before and after placement of 4 different silicone plugs (n=4 samples, Figure 3b). Silicone earplugs were found to provide a mild to moderate temporary hearing loss, with attenuation levels ranging from <10 dB at frequencies less than 4 kHz (mean: 10±7.5 dB) toand 10-35 dB at frequencies greater than 4 kHz (mean: 23.7±9.4 dB; Figure 2b). The frequency-specific attenuation levels that the silicone plugs provide are consistent with that of foam ear plugs in other species (*barn owl*: Knudsen et al., 1984; *chinchilla*: Lupo et al., 2011; *ferret*: Moore et al., 1999).

#### Startle-based method for assessing spatial hearing

Experiments were performed in a double-walled, sound-attenuating chamber (interior dimensions: ∼3 × 3 × 3 m; Industrial Acoustics Company, IAC, Bronx, NY) lined with acoustical foam. The animal was placed in a custom-built acoustically transparent wire-mesh cage mounted on a polyvinyl chloride (PVC) post anchored to a flexible polycarbonate platform. The cage was oriented such that the animal faced forward towards the center loudspeaker (Figure 6a). Proper orientation of the animal was visually monitored using a closed-circuit infrared camera. Prepulse stimuli were presented from 25 identical loudspeakers (Morel MDT-20) spaced along a 1m radius semicircular hoop at 7.5° increments, from −90° (right) to +90° (left). The array of speakers was oriented horizontally (i.e. 0° elevation) for all experiments. An additional loudspeaker (Faital Pro HF102) positioned ∼25 cm above the animal provided the startle-eliciting stimuli (20ms duration broadband noise bursts presented at110-120 dB SPL). The startle response of the animal was captured using a cage-mounted accelerometer (Analog Devices ADXL335). Generation of the sound stimuli and recording of the startle signal was controlled by custom written MATLAB (MathWorks) software controlling three Tucker-Davis Technologies (TDT) Real-time Processors (RP2.1). The startle response amplitude was calculated as the RMS of the accelerometer output in the first 100ms period after the delivery of the ES.

Behavioral spatial acuity was assessed using the speaker-swap (SSwap) paradigm (Allen and Ison, 2010; Greene et al., 2018), which quantifies the smallest angle for which a change in speaker location can be detected by the animal (i.e., the minimum audible angle; cf. Mills 1958). Guinea pigs reflexively startle in response to loud, unexpected sounds (Figure 6b), but presentation of a detectable cue (the “prepulse”) prior to the loud sound reduces the startle response (prepulse inhibition, PPI; Figure 6c). Here, the prepulse signal is a swap of continuous noise (broadband or high-pass) presented from one speaker to a second speaker, prior to presentation of a startle stimulus (interstimulus interval, ISI, between prepulse and ES: 300 ms). The direction of the swap was also randomized (e.g. noise swaps from *left-to-right* or *right-to-left* speakers). Five angles of the speaker swap prepulse were tested (7.5 °, 15 °, 30 °, 45 °, 90 °; Figure 6a).

Responses are the average of at least 10 repetitions, and are shown in units of prepulse inhibition (PPI). PPI is defined as 1 minus the ratio of RMS amplitude of the prepulse condition (speaker swap present) divided by the control condition (no speaker swap):

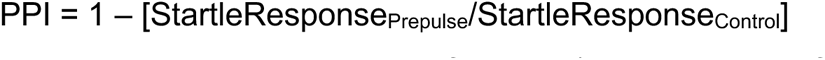

A PPI of “1” indicates a complete suppression of startle (i.e., detection of the swap), a “0” indicates that the startle response for the swap condition is no different from the control response (i.e., no detection of swap), and “<0” indicates a facilitation, or enhancement, of the startle response. The minimum audible angle is defined as the smallest speaker swap angle that the animal reliably detected, indicated by a cross-trial averaged startle response significantly different from that observed in the absence of the aprepulse (swap) stimulus. Significance testing was completed using a repeated-measures analysis of variance (ANOVA) to determine whether prepulse conditions were significantly different from no-prepulse conditions (controls), followed by Bonferroni-corrected two-tailed Student’s t-tests to determine which angles were significantly different from control angles (PPI=0).

#### Single-unit extracellular recordings in the auditory midbrain

After completion of ABR and behavioral tests, anesthetized animals were placed in a double-walled sound attenuating chamber (IAC, Bronx, NY) for physiological testing. Following methods of Jones et al. (2015), hair was removed from the ventral aspect of the neck and a tracheal cannula was implanted to monitor respiration and ensure ease of breathing. The animal was then placed on a custom bite bar fixed to a stereotaxic apparatus (model 1430, David Kopf Instruments). Hair was removed from the top of the head, and a midline incision was made to expose the skull and the bony ear canals. Custom stainless-steel earpieces were then secured into each ear canal. Closed-field speakers were attached to the left and right earpieces to present acoustic stimuli. Probe microphones (Type 4182, Bruel and Kjaer) incorporated in the insert earpieces were used to calibrate tones between 50Hz-20kHz. A ∼2-3 mm craniotomy was made near the midline of the skull (at the interaural axis) to expose cortex overlying the inferior colliculus (IC). In some experiments, the overlying cortex was aspirated to visualize the inferior colliculi, confirming the correct placement of the electrode. For the “early CHL” animals, recordings were made in the IC both contralateral and ipsilateral to the previously-plugged ear (Figure 7c). Parylene-coated Tungsten microelectrodes (2-5 MΩ; MicroProbe, Inc) were secured to a micromanipulator and were advanced remotely outside of the chamber while presenting search stimuli (repeating tone sweeps, from ∼100Hz-20kHz). The correct placement of the electrode is validated when neurons reliably respond to auditory stimuli. Neurons in the central nucleus of the inferior colliculus (ICC) were targeted for physiological assessments. The ICC is a tonotopically organized auditory midbrain region where neurons are responsive to low frequencies in the dorsal ICC (<3 kHz in the guinea pig) and higher frequencies in ventral ICC (>16 kHz; Dong et al., 2010). Thus, as the electrode is advanced ventrally, the frequency selectivity of ICC neurons increases (i.e., from ∼100Hz to ∼20kHz). ICC unit responses were amplified (ISO-80, WPI, Sarasota, FL; Stanford Research Systems, SRS 560, Sunnyvale, CA) and filtered (300-3000 Hz). A BAK amplitude-time window discriminator (Model DDIS-1, Mount Airy, MD) was used to evaluate neuron responses and spike times were stored at a precision of 1 μs via a Tucker-Davis Technologies (TDT, Alachua, FL) RV8.

Units were selected for further study if the spike waveforms exhibited good signal-to-noise ratio, the responses were strong and non-adapting to auditory stimuli, and the responses exhibited dorso-ventral tonotopy. For each isolated ICC neuron, the characteristic frequency (CF), rate-level function at CF, and binaural sensitivity to interaural level differences (ILDs) was assessed. The CF for each isolated unit was estimated from obtaining a three-dimensional plot of signal frequency (kHz), signal intensity (dB), and firing rate (spikes/sec; Supplemental Figure 4a). After determining the CF of an isolated unit, the threshold of the neuron was assessed by varying the intensity of a CF tone presented to the ear contralateral to the electrode (e.g., recording in the left IC, the intensity presented to the right ear would be varied) to get a rate-level function (Supplemental Figure 5a). From the rate-level function, the contralateral signal intensity that elicited approximately 50% of the maximal firing rate (typically 50-60 dB SPL and ∼20 dB above threshold) was documented and used for subsequent binaural testing.

Binaural sensitivity to ILDs was assessed by fixing the sound intensity of a 200-ms CF tone at the contralateral ear (the “excitatory” ear) at ∼20 dB above threshold and varying the sound intensity presented at the ipsilateral ear (the “inhibitory” ear) across a ±30 dB range in 5 dB steps; Figure 7a-b, allowing us to manipulate the level of inhibition with a constant level of excitation. For example, when recording in the left IC, if the threshold of an isolated unit was 40 dB SPL, then the level at the right ear was be fixed at 60 dB SPL and the intensity presented to the left ear varied from 30 to 90 dB SPL. Each ILD cue was presented 20 times, with cue values presented in random order over the course of a testing block. Raw rate-ILD data were fit with a four-parameter sigmoidal logistic function of the form, y(*α*) = *A* + ((*D*-*A*)/(1+*expB*(*α*-*C*))) where *α* is the ILD (ipsi SPL ─ contra SPL), *A* is the minimum firing rate, *D* is the maximum firing rate, *C* is the inflection point (referred to as the “50% inflection point”), and *B* is a width parameter, the sign of which also controls the direction of inflection. Units were classified as ILD-sensitive if the firing rate was modulated by at least 50% with varying sound intensity to the ipsilateral (inhibitory) ear and the R^2^ of the fit was at least 0.70. The fit parameters include the 50% inflection point (i.e., half-maximum ILD value, or the ILD yielding 50% of maximal firing), the rate-ILD slope (spikes/s/dB ILD, computed at half-max ILD), and the ILD dynamic range (the range of ILD producing 10-90% of maximal firing; Supplemental Figure 3a; Tollin et al., 2008; Tsai et al., 2010). A total of 107 isolated units were classified as ILD-sensitive according to these criteria, including n=40 units from control animals and n=67 units from CHL animals (of these, 39 were recorded in the IC ipsilateral to the earplug, and 29 were recorded in the IC contralateral to the earplug.

The ILD coding acuity of ICC neurons was next assessed according to the Fisher information (FI) conveyed by fitted rate-ILD functions. FI can be used to quantify the precision with which a neuron’s responses discriminate adjacent stimulus values (e.g. two ILDs). Here, FI is expressed in physical units (1/dB^2^) which can be used to compute neural ILD discrimination (i.e., ILD just-noticeable-differences, JNDs), and allow for a comparison between neural and behavioral acuity of sound source locations. The FI computation used here is also described in detail elsewhere (Seung and Sompolinsky, 1993; Tollin et al., 2008; Jones et al., 2015; Brown and Tollin, 2016; Benichoux et al., 2017). FI is computed as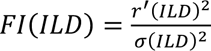

Where r’(ILD) is the derivative of the spike count vs ILD function with respect to ILD, and σ(ILD)^2^ is the variance of spike count across ILD. Here, we assumed a Poisson distribution such that the numerical value of σ(ILD)^2^ was set to equal the numerical value of the spike count (i.e., the number of spikes expected for a 1 second stimulus). Since FI considers the slope and variance of spiking across stimulus values, it can easily be converted to a signal detection theoretic measure (Green and Swets, 1966; Brown and Tollin, 2016) by evaluating the change in stimulus value (ILD) necessary to achieve 1 standard separation in spiking distributions (i.e., the neural JND for a d’ of 1, which corresponds to 76% correct discrimination in a 2-alternative forced choice psychophysical task). This value is given by the following:

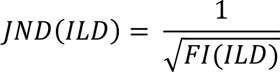

where *JND(ILD)* gives the neuron’s ILD JND, and *FI(ILD)* is the FI as a function of ILD as computed in the *FI(ILD)* equation above. See Figure 7d for an overview of the FI computation pipeline.

## Abbreviations

(ABR): auditory brainstem response
(BIC): binaural interaction component
(CHL): conductive hearing loss
(IC): inferior colliculus
(ILD): interaural level difference
(ITD): interaural time difference
(LSO): lateral superior olive

## Supplemental Figures

**Supplemental Figure 1.**
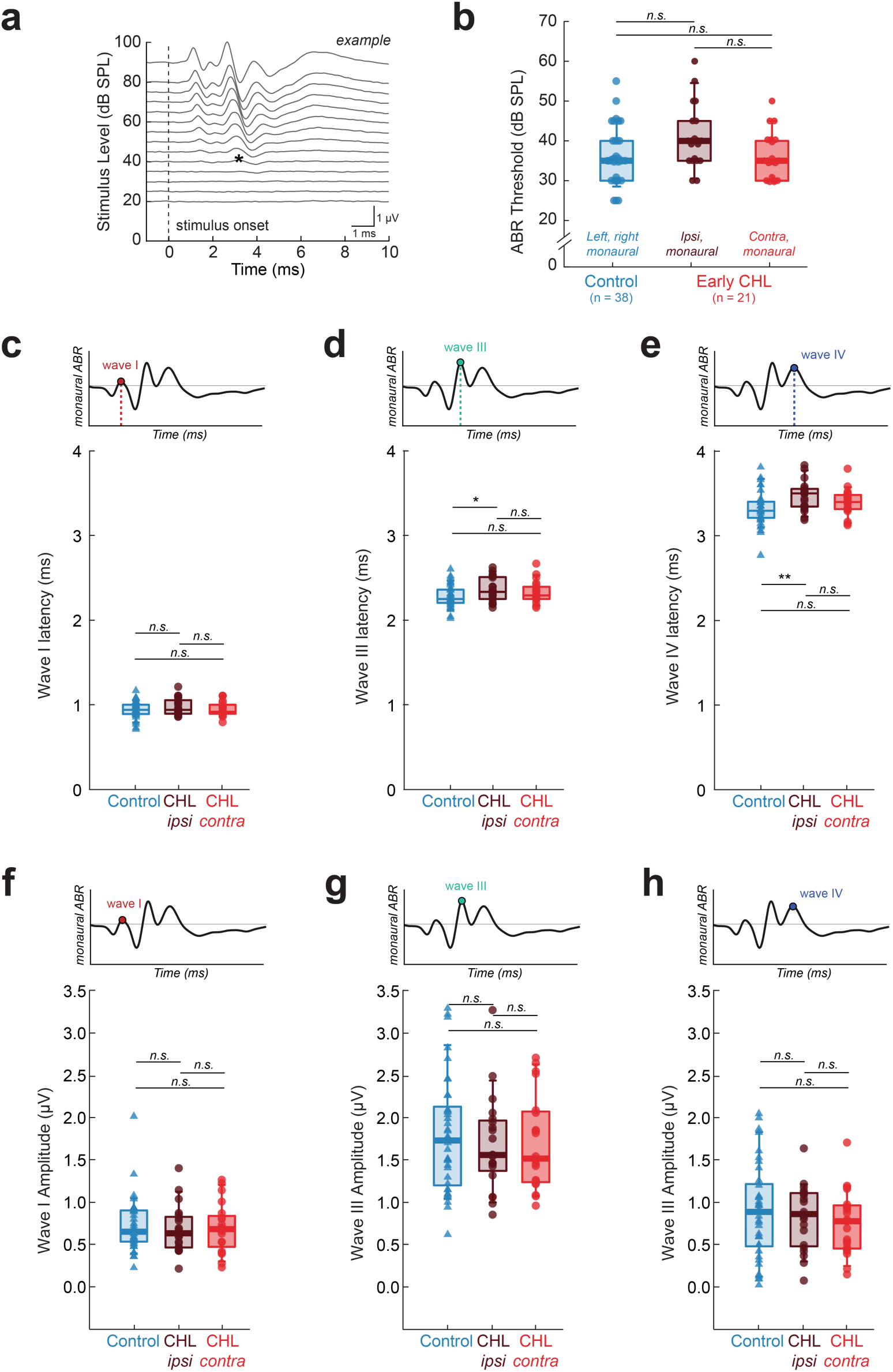
Developmental CHL does not alter click-evoked monaural ABRs. **(a)** Example left monaural ABR trace for a littermate control animal (#124111). Traces are in response to broadband click stimuli and are shown for different stimulus intensities (20-90 dB SPL in 5 dB steps). Traces shown are the average of 500 repetitions. The dotted vertical line indicates the stimulus onset of the click. ABR thresholds were measured by identifying the lowest stimulus level at which a response was visually detectable, regardless of the specific wave. For instance, the threshold for this example would be 40 dB SPL (black asterisk). (**b**) Click-ABR thresholds are not significantly elevated in animals raised with CHL (Mann-Whitney U, n.s.: p>0.01). Circles depict identified thresholds from monaural ABRs of normal animals (black, left and right ABRs were combined) and animals raised with an earplug (red). For the Early CHL animals, data were split into two groups: thresholds ipsilateral (dark red) and contralateral (bright red) to the previously occluded ear. Boxplots indicating the median, 10^th^, 25^th^, 75^th^, and 90^th^ percentiles with error bars are shown for each group. The median thresholds (± SD) for controls, Early CHL_ipsi_, and Early CHL_contra_ are 35±7.2 dB SPL, 40±7.9 dB SPL, and 35±5.8 dB SPL, respectively. (**c-e**) Latencies (ms) for monaural ABR waves I (**a**), III (**b**), and IV (**c**) for controls (black) and early CHL animals (ipsi to CHL: dark red; contra to CHL: bright red). Wave II was not included as it is not always identifiable in the ABR traces. (**f-h**) Amplitudes (µV) for monaural ABR waves I (**f**), III (**g**), and IV (**h**) for control and early CHL animals. For (**c-h**), the top inset shows an example ABR with the wave of focus emphasized. Each symbol indicates measurements taken from the left and right ear ABRs; controls include both the left and right (n=38 traces), whereas the CHL groups are split into measurements ipsilateral and contralateral to the developmental CHL (n=21 traces for both groups). Boxplots indicating the median, 10^th^, 25^th^, 75^th^, and 90^th^ percentiles with error bars are shown for each group. n.s. (Mann-Whitney U): p>0.05.

**Supplemental Figure 2.**
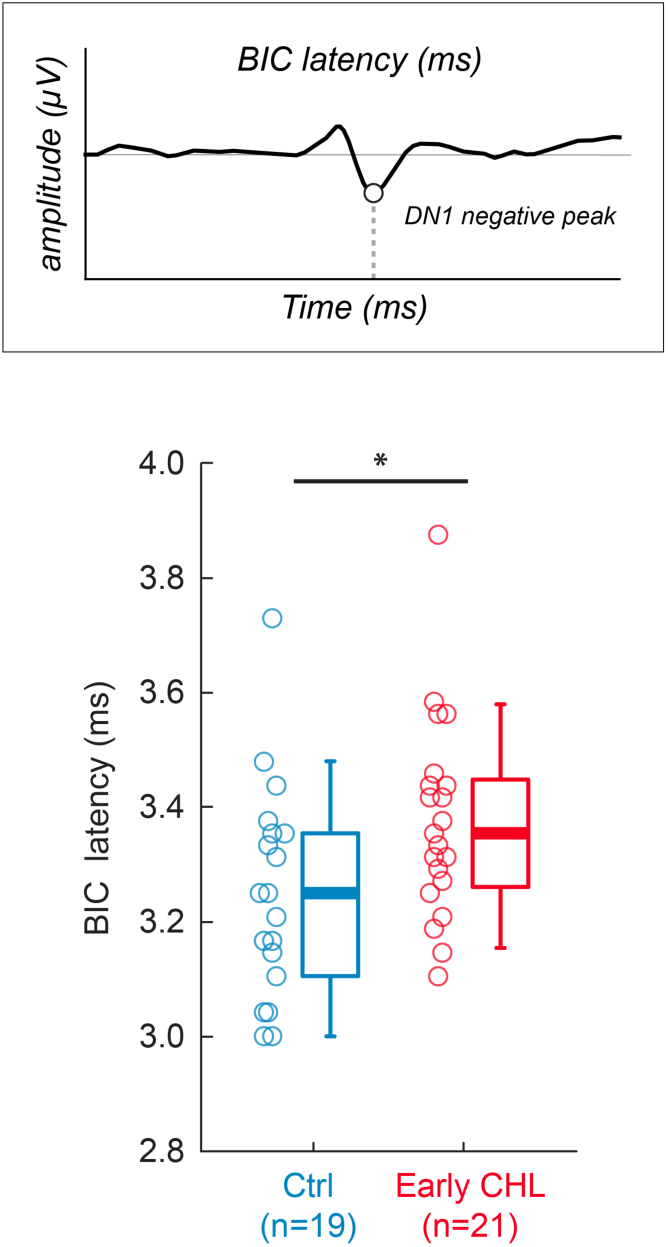
Early CHL alters BIC latency. Top panel: The latency (ms) of the BIC is defined as the time from stimulus onset to the time of the DN1 negative peak of the BIC. Bottom panel: BIC latency for Control (n=19) and Early CHL (n=21) animals. Circles indicate individual values and the boxplots show the group median and the 10^th^, 25^th^, 75^th^, and 90^th^ percentiles along with the associated error bars. Early CHL animals exhibit longer BIC latency than their littermate controls (Ctrl median±SD: 3.25±0.19 ms; Early CHL: 3.38±0.18 ms; Mann-Whitney U, p=0.03).

**Supplemental Figure 3.**
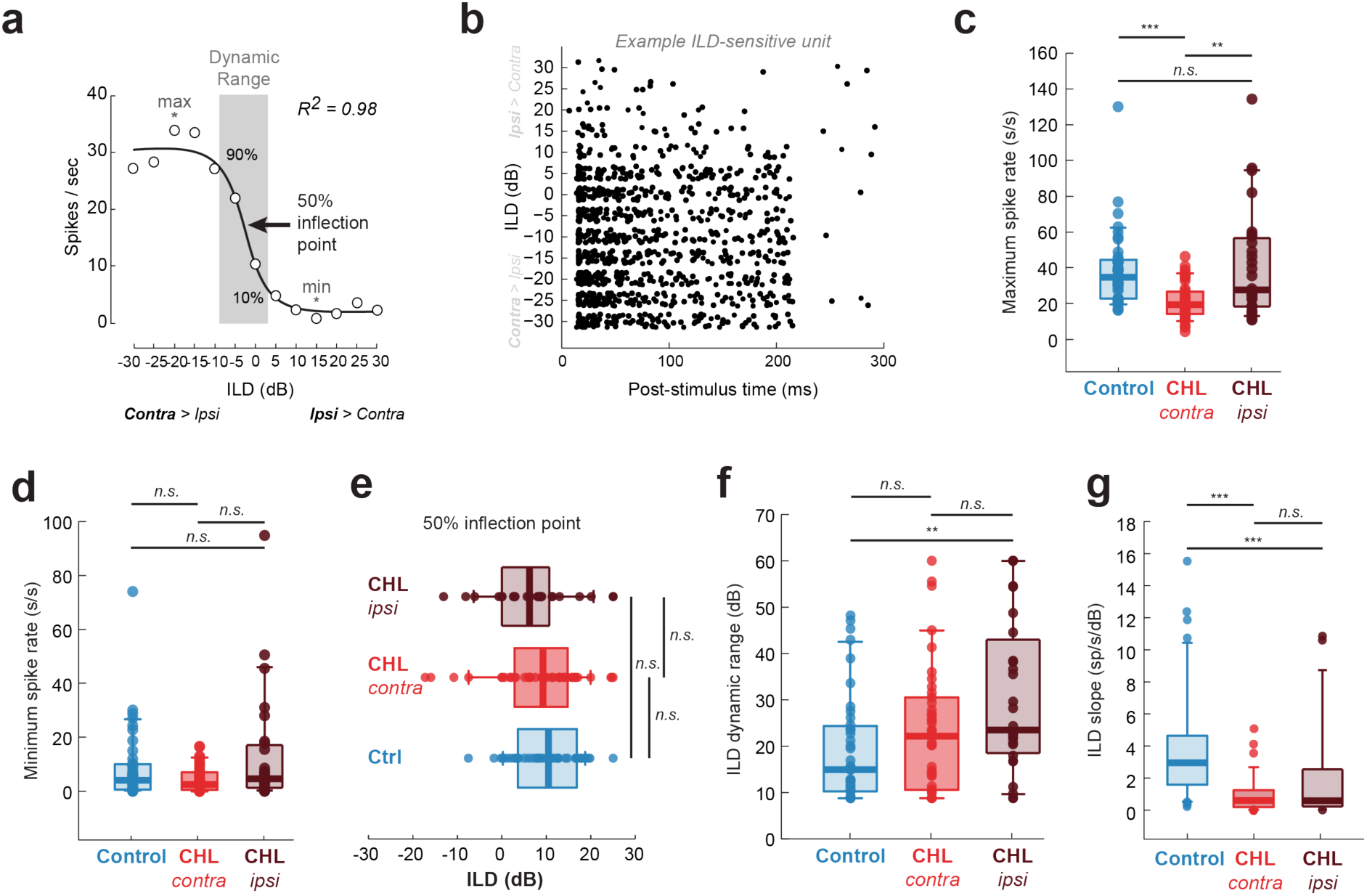
Rate-ILD functions for control and Early CHL IC neurons. (**a**) Example rate-ILD function for an ILD-sensitive IC neuron. A sigmoid function is fit to all data to compare parameters between groups. Neurons were considered ILD-sensitive when their discharge rates were modulated by ≥ 50% with increasing sound intensity to the ipsilateral ear (i.e., increasing the inhibitory drive). (**b**) Spike rasters for example ILD-sensitive unit shown in (**a**). From the rate-ILD functions, maximum (**c**) and minimum (**d**) spike rates were plotted from controls (blue) and early CHL animals (contra to plug: bright red; ipsi to plug: dark red). Maximum spike rates from neurons contralateral to the previous CHL are significantly smaller than from control neurons and neurons ipsilateral to previous CHL. There are no significant differences between the groups for minimum spike rates. (**e**) The 50% inflection point from the sigmoid fits were not significantly different between groups. (**f**) The ILD dynamic range for neurons ipsilateral to the previous CHL were significantly different from controls, whereas they were not significant for neurons contralateral to the previous CHL. (**g**) The ILD slope was significantly smaller for both CHL groups compared to controls. n.s.: p>0.05; *: p<0.05; **: p<0.01; ***: p<0.001.

**Supplemental Figure 4.**
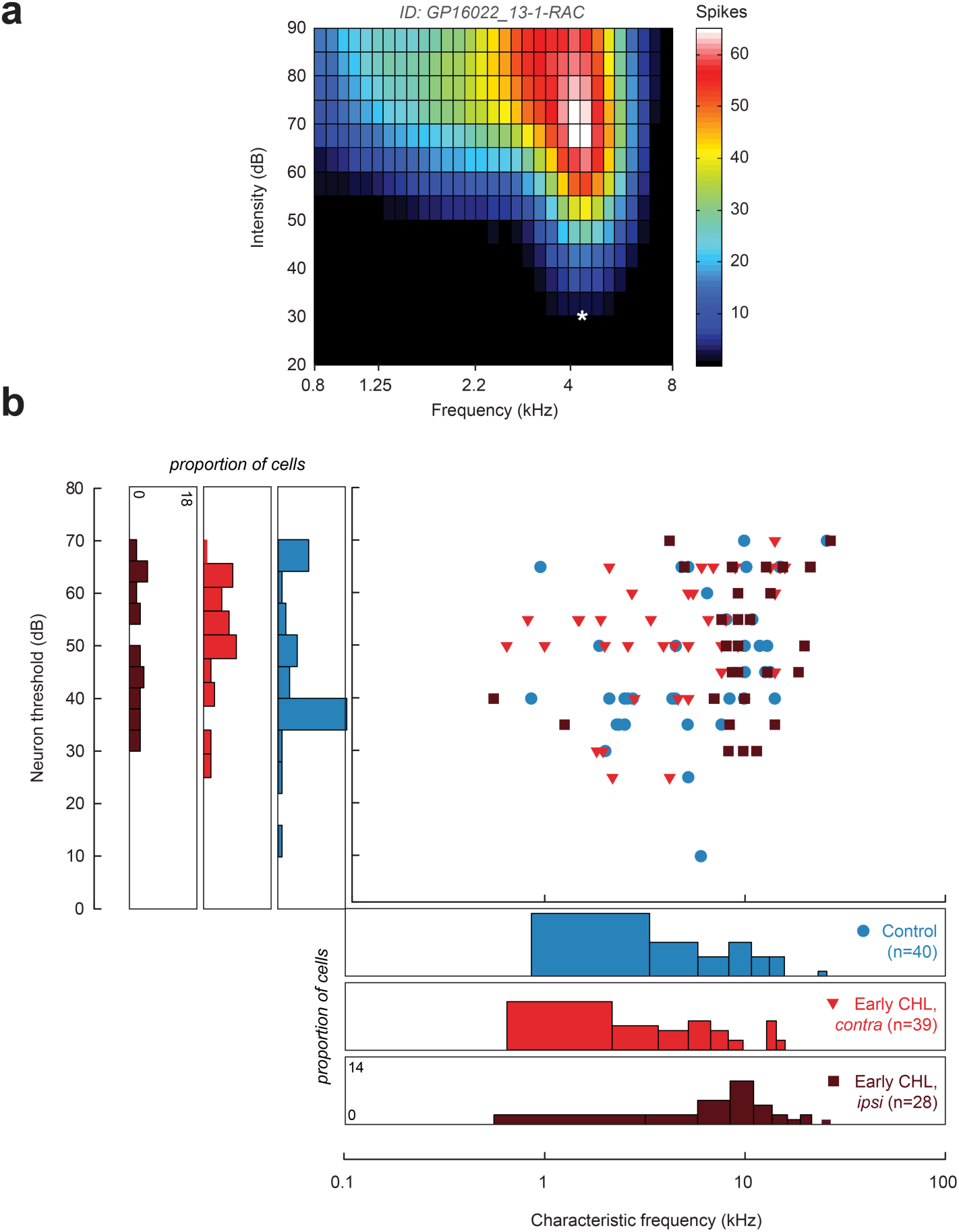
Monaural inferior colliculus (IC) tuning properties from control and Early CHL animals. (**a**) Response area curves (RACs) were collected from all IC neurons to determine the characteristic frequency (CF) and the unit threshold. (**a**) Example RAC for an IC neuron. The asterisk (*) indicates the frequency at which the neuron is most sensitive to (CF: 4.7 kHz; threshold: 35 dB). (**b**) Neuron threshold as a function of CF for controls (blue) and from early CHL animals (neurons contralateral to CHL, bright red; neurons ipsilateral to CHL, dark red). Only neurons that are sensitive to interaural level differences (ILD) are shown here (see Methods for inclusion criteria). Histograms for CF and threshold are plotted on the x- and y-axes, respectively, to show distribution of CF and thresholds for control and Early CHL neurons.

**Supplemental Figure 5.**
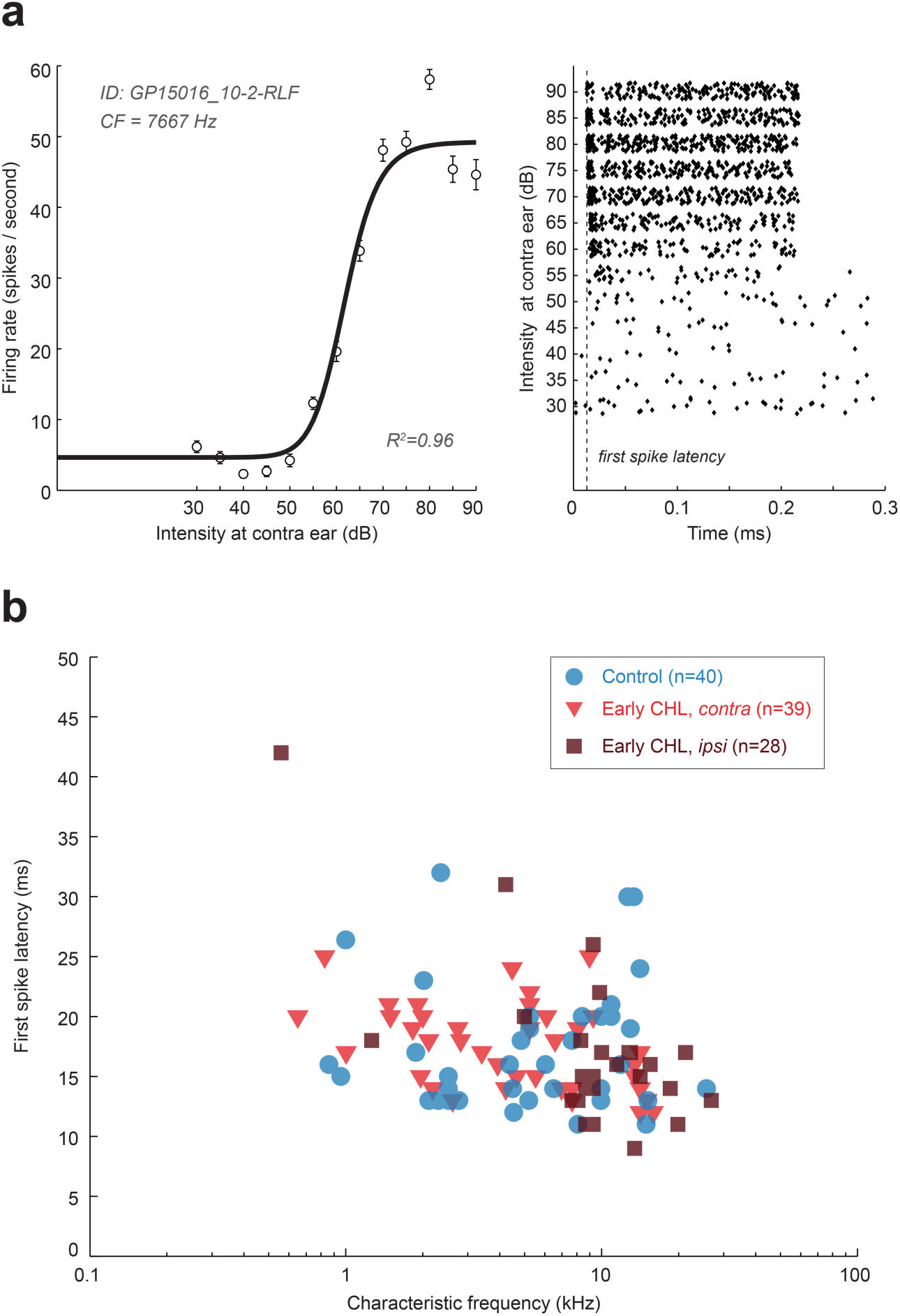
Rate level functions and first spike latencies for IC neurons. **(a)** Rate level functions (RLFs) were collected from IC to assess unit thresholds and first spike latencies (i.e., monaural response properties). The left panel shows an example RLF for an IC neuron. Frequency tones at the CF of the unit are presented at varying intensities at the contralateral ear (e.g., recording in the left IC, present tones from the right ear speaker). The right panel shows the spike raster for each intensity as function of time for the example neuron. The dotted line indicates the first spike latency (ms). (**b**) First spike latency as a function of CF from controls (blue) from early CHL animals (neurons contralateral to CHL, bright red; neurons ipsilateral to CHL, dark red). There are no significant differences in latencies between Control or Early CHL neurons (F_2,104_ = 0.03, p=0.98).

## References

A. Bf (2002) Presbyacusis and the Auditory Brainstem Response. J Speech, Lang Hear Res 45:1249–1261 Available at: 10.1044/1092-4388(2002/100).

Allen CB, Celikel T, Feldman DE (2003) Long-term depression induced by sensory deprivation during cortical map plasticity in vivo. Nat Neurosci 6:291.

changes in azimuthal sound location: angular separation, spectral composition, and sound level. Behav Neurosci 124:265–277

Anbuhl KL, Benichoux V, Greene NT, Brown AD, Tollin DJ (2017) Development of the head, pinnae, and acoustical cues to sound location in a precocial species, the guinea pig (Cavia porcellus). Hear Res 356.

Beiderbeck B, Myoga MH, Müller NIC, Callan AR, Friauf E, Grothe B, Pecka M (2018) Precisely timed inhibition facilitates action potential firing for spatial coding in the auditory brainstem. Nat Commun 9:1–13.

Bender KJ (2006) Synaptic Basis for Whisker Deprivation-Induced Synaptic Depression in Rat Somatosensory Cortex. J Neurosci 26:4155–4165.

Benichoux V, Brown AD, Anbuhl KL, Tollin DJ (2017) Representation of multidimensional stimuli: Quantifying the most informative stimulus dimension from neural responses. J Neurosci 37.

Benichoux V, Ferber A, Hunt S, Hughes E, Tollin D (2018) Across Species “Natural Ablation” Reveals the Brainstem Source of a Noninvasive Biomarker of Binaural Hearing. J Neurosci 38:8563–8573.

Bennett KE, Haggard MP, Silva PA, Stewart IA (2001) Behaviour and developmental effects of otitis media with effusion into the teens. Arch Dis Child 85:91–95.

Beutelmann R, Laumen G, Tollin D, Klump GM (2015) Amplitude and phase equalization of stimuli for click evoked auditory brainstem responses. J Acoust Soc Am 137:EL71–EL77.

Blake R, Hirsch H V (1975) Deficits in binocular depth perception in cats after alternating monocular deprivation. Science (80-) 190:1114 LP–1116.

Brown AD, Tollin DJ (2016) Slow Temporal Integration Enables Robust Neural Coding and Perception of a Cue to Sound Source Location. J Neurosci 36:9908–9921.

Brugge JF, Orman SS, Coleman JR, Chan JCK, Phillips DP (1985) Binaural interactions in cortical area AI of cats reared with unilateral atresia of the external ear canal. Hear Res 20:275–287.

Carney LH, Sarkar S, Abrams KS, Idrobo F (2011) Sound-localization ability of the Mongolian gerbil (Meriones unguiculatus) in a task with a simplified response map. Hear Res 275:89– 95.

Clifton R, Gwiazda J, Bauer JA, Clarkson MG, Held RM (1988) Growth in head size during infancy: Implications for sound localization. Dev Psychol 24:477–483.

Clopton BM, Silverman MS (1977) Plasticity of binaural interaction. II. Critical period and changes in midline response. J Neurophysiol 40:1275–1280.

Coleman JE, Nahmani M, Gavornik JP, Haslinger R, Heynen AJ, Erisir A, Bear MF (2010) Rapid Structural Remodeling of Thalamocortical Synapses Parallels Experience-Dependent Functional Plasticity in Mouse Primary Visual Cortex. J Neurosci 30:9670– 9682.

Delb W, Strauss DJ, Hohenberg G, Plinkert PK, Delb W (2003) The binaural interaction component (BIC) in children with central auditory processing disorders (CAPD): El componente de interactión binaural (BIC) en niños con desórdenes del procesamiento central auditivo (CAPD). Int J Audiol 42:401–412.

Dews PB, Wiesel TN (1970) Consequences of monocular deprivation on visual behaviour in kittens. J Physiol 206:437–455.

Dong S, Rodger J, Mulders WH A M, Robertson D (2010) Tonotopic changes in GABA receptor expression in guinea pig inferior colliculus after partial unilateral hearing loss. Brain Res 1342:24–32.

Dum N (1984) Postnatal development of the auditory evoked brainstem potentials in the guinea pig. Acta Otolaryngol 97:63–68.

Eggermont JJ, Moore JK (2012) Morphological and Functional Development of the Auditory Nervous System. In, pp 61–105. Springer New York.

Espinosa JS, Stryker MP (2012) Development and Plasticity of the Primary Visual Cortex. Neuron 75:230–249.

Ferber AT, Benichoux V, Tollin DJ (2016) Test-retest reliability of the binaural interaction component of the auditory brainstem response. Ear Hear 37:e291.

Fields RD (2015) A new mechanism of nervous system plasticity: activity-dependent myelination. Nat Rev Neurosci 16:756–767.

Folsom RC, Weber BA, Thompson G (1983) Auditory brainstem responses in children with early recurrent middle ear disease. Ann Otol Rhinol Laryngol 92:249–253.

Fox K (1992) A critical period for experience-dependent synaptic plasticity in rat barrel cortex. J Neurosci 12:1826–1838.

Franken TP, Joris PX, Smith PH (2018) Principal cells of the brainstem’s interaural sound level detector are temporal differentiators rather than integrators. Elife 7:e33854.

Furst M, Eyal S, Korczyn AD (1990) Prediction of binaural click lateralization by brainstem auditory evoked potentials. Hear Res 49:347–359.

Green DM, Swets JA (1966) Signal detection theory and psychophysics. New York-London-Sydney: John WiIey & Sons.

Greene NT, Anbuhl KL, Ferber AT, DeGuzman M, Allen PD, Tollin DJ (2018) Spatial hearing ability of the pigmented Guinea pig (Cavia porcellus): Minimum audible angle and spatial release from masking in azimuth. Hear Res.

Greene NT, Anbuhl KL, Williams W, Tollin DJ (2014) The acoustical cues to sound location in the guinea pig (Cavia porcellus). Hear Res 316C:1–15.

Gunnarson AD, Finitzo T (1991) Conductive hearing loss during infancy: Effects on later auditory brain stem electrophysiology. J Speech, Lang Hear Res 34:1207–1215.

Hall III JW, Grose JH (1994) Development of temporal resolution in children as measured by the temporal modulation transfer function. J Acoust Soc Am 96:150–154.

Hall JW, Grose JH (1993) The effect of otitis media with effusion on the masking-level difference and the auditory brainstem response. J Speech, Lang Hear Res 36:210–217.

Hall JW, Grose JH, Pillsbury HC (1995) Long-term effects of chronic otitis media on binaural hearing in children. Arch Otolaryngol Neck Surg 121:847–852.

Hartley DEH, Moore DR (2003) Effects of conductive hearing loss on temporal aspects of sound transmission through the ear. Hear Res 177:53–60.

Heffner R, Heffner H, Masterton B (1971) Behavioral Measurements of Absolute and Frequency-Difference Thresholds in Guinea Pig. J Acoust Soc Am 49:1888–1895.

Hirsch JA, Oertel D (1988) Intrinsic properties of neurones in the dorsal cochlear nucleus of mice, in vitro. J Physiol 396:535–548.

Hofer SB, Mrsic-Flogel TD, Bonhoeffer T, Hübener M (2009) Experience leaves a lasting structural trace in cortical circuits. Nature 457:313–317.

Hubel DH, Wiesel TN (1970) The period of susceptibility to the physiological effects of unilateral eye closure in kittens. J Physiol 206:419–436.

Hurley RM, Hurley A (1995) The auditory brainstem response in children with histories of otitis media. In: Seminars in Hearing, pp 37–42. Copyright© 1995 by Thieme Medical Publishers, Inc.

Irvine DR, Gago G (1990) Binaural interaction in high-frequency neurons in inferior colliculus of the cat: effects of variations in sound pressure level on sensitivity to interaural intensity differences. J Neurophysiol 63:570–591.

Ito T, Tokuriki M, Shibamori Y, Saito T, Nojyo Y (2004) Cochlear nerve demyelination causes prolongation of wave I latency in ABR of the myelin deficient (md) rat. Hear Res 191:119– 124.

Jewett DL (1970) Volume-conducted potentials in response to auditory stimuli as detected by averaging in the cat. Electroencephalogr Clin Neurophysiol 28:609–618.

Jones HG, Brown AD, Koka K, Thornton JL, Tollin DJ (2015) Sound frequency-invariant neural coding of a frequency-dependent cue to sound source location. J Neurophysiol 114:531– 539.

Jonson KM, Lyle JG, Edwards MJ, Penny RHC (1975) Problems in behavioural research with the guinea pig: A selective review. Anim Behav 23:632–639.

Kandler K, Clause A, Noh J (2009) Tonotopic reorganization of developing auditory brainstem circuits. Nat Neurosci 12:711.

Keating P, Dahmen JC, King AJ (2013a) Context-Specific Reweighting of Auditory Spatial Cues following Altered Experience during Development. Curr Biol 23:1291–1299.

Keating P, Dahmen JC, King AJ (2015) Complementary adaptive processes contribute to the developmental plasticity of spatial hearing. Nat Neurosci 18:185–187.

Keating P, Nodal FR, Gananandan K, Schulz AL, King AJ (2013b) Behavioral Sensitivity to Broadband Binaural Localization Cues in the Ferret. J Assoc Res Otolaryngol 14:561–572.

Kim G, Kandler K (2010) Synaptic changes underlying the strengthening of GABA/glycinergic connections in the developing lateral superior olive. Neuroscience 171:924–933.

Knudsen EI, Esterly SD, Knudsen PF (1984) Monaural occlusion alters sound localization during a sensitive period in the barn owl. J Neurosci 4:1001–1011.

Koiek, S., Brandt, C., Schmidt, J. H., & Neher, T. (2022). Monaural and binaural phase sensitivity in school-age children with early-childhood otitis media. International Journal of Audiology, 61(12), 1054–1061.

Kumpik DP, King AJ (2018) A review of the effects of unilateral hearing loss on spatial hearing. Hear Res.

Laska M, Walger M, Schneider I, von Wedel H (1992) Maturation of binaural interaction components in auditory brainstem responses of young guinea pigs with monaural or binaural conductive hearing loss. Eur Arch Oto-Rhino-Laryngology 249:325–328.

Laumen G, Ferber AT, Klump GM, Tollin DJ (2016) The Physiological Basis and Clinical Use of the Binaural Interaction Component of the Auditory Brainstem Response. Ear Hear 37:e276–e290.

Ludwig AA, Zeug M, Schönwiesner M, Fuchs M, Meuret S (2019) Auditory localization accuracy and auditory spatial discrimination in children with auditory processing disorders. Hear Res 377:282–291.

Lupo JE, Koka K, Thornton JL, Tollin DJ (2011) The effects of experimentally induced conductive hearing loss on spectral and temporal aspects of sound transmission through the ear. Hear Res 272:30–41.

Maffei A, Nelson SB, Turrigiano GG (2004) Selective reconfiguration of layer 4 visual cortical circuitry by visual deprivation. Nat Neurosci 7:1353–1359.

Miller GL, Knudsen EI (2003) Adaptive plasticity in the auditory thalamus of juvenile barn owls. J Neurosci 23:1059–1065.

Mishra, S. K., & Moore, D. R. (2023). Auditory deprivation during development alters efferent neural feedback and perception. Journal of Neuroscience, 43(25), 4642–4649.

Mogdans J, Knudsen EI (1993) Early monaural occlusion alters the neural map of interaural level differences in the inferior colliculus of the barn owl. Brain Res 619:29–38.

Moore DR, Hartley DEH, Hogan SCM (2003) Effects of otitis media with effusion (OME) on central auditory function. Int J Pediatr Otorhinolaryngol 67:S63–S67.

Moore DR, Hine JE, Jiang ZD, Matsuda H, Parsons CH, King AJ (1999) Conductive Hearing Loss Produces a Reversible Binaural Hearing Impairment. 19:8704–8711.

Nittrouer, S., & Lowenstein, J. H. (2024). Early otitis media puts children at risk for later auditory and language deficits. International Journal of Pediatric Otorhinolaryngology, 176, 111801.

Polley DB, Thompson JH, Guo W (2013) Brief hearing loss disrupts binaural integration during two early critical periods of auditory cortex development. Nat Commun 4:2547.

Popescu M V., Polley DB (2010) Monaural Deprivation Disrupts Development of Binaural Selectivity in Auditory Midbrain and Cortex. Neuron 65:718–731.

Prosen C a, Petersen MR, Moody DB, Stebbins WC (1978) Auditory thresholds and kanamycin-induced hearing loss in the guinea pig assessed by a positive reinforcement procedure. J Acoust Soc Am 63:559–566.

Reichman J, Healey WC (1983) Learning disabilities and conductive hearing loss involving otitis media. J Learn Disabil 16:272–278.

Rema V, Armstrong-James M, Ebner FF (2003) Experience-Dependent Plasticity Is Impaired in Adult Rat Barrel Cortex after Whiskers Are Unused in Early Postnatal Life. J Neurosci 23:358 LP – 366.

Riedel H, Kollmeier B (2006) Interaural delay-dependent changes in the binaural difference potential of the human auditory brain stem response. Hear Res 218:5–19.

Sanes DH (1990a) An in vitro analysis of sound localization mechanisms in the gerbil lateral superior olive. J Neurosci 10:3494–3506.

Sanes DH (1990b) An in vitro analysis of sound localization mechanisms in the gerbil lateral superior olive. J Neurosci 10:3494–3506.

Sanes DH, Rubel EW (1988) The ontogeny of inhibition and excitation in the gerbil lateral superior olive. J Neurosci 8:682–700.

Sanes DH, Takács C (1993) Activity-dependent refinement of inhibitory connections. Eur J Neurosci 5:570–574.

Seung HS, Sompolinsky H (1993) Simple models for reading neuronal population codes. Proc Natl Acad Sci 90:10749–10753.

Shepherd GMG, Pologruto TA, Svoboda K (2003) Circuit Analysis of Experience-Dependent Plasticity in the Developing Rat Barrel Cortex. Neuron 38:277–289.

Silverman MS, Clopton BM (1977) Plasticity of binaural interaction. I. Effect of early auditory deprivation. J Neurophysiol 40:1266–1274.

Thornton JL, Chevallier KM, Koka K, Lupo JE, Tollin DJ (2012) The conductive hearing loss due to an experimentally induced middle ear effusion alters the interaural level and time difference cues to sound location. J Assoc Res Otolaryngol 13:641–654.

Timney B, Mitchell DE, Cynader M (1980) Behavioral evidence for prolonged sensitivity to effects of monocular deprivation in dark-reared cats. J Neurophysiol 43:1041–1054.

Tollin D, Blumberg M (2010) Development of sound localization mechanisms. Oxford Handb Dev Behav Neurosci:262–282.

Tollin DJ (2003) The Lateral Superior Olive: A Functional Role in Sound Source Localization. Neurosci 9:127–143.

Tollin DJ, Koka K, Tsai JJ (2008) Interaural level difference discrimination thresholds for single neurons in the lateral superior olive. J Neurosci 28:4848–4860.

Tolnai S, Klump GM (2020) Evidence for the origin of the binaural interaction component of the auditory brainstem response. Eur J Neurosci 51:598–610.

Tsai JJ, Koka K, Tollin DJ (2010) Varying overall sound intensity to the two ears impacts interaural level difference discrimination thresholds by single neurons in the lateral superior olive. J Neurophysiol 103:875–886.

Ungan P, Yaǧcioǧlu S, Özmen B (1997) Interaural delay-dependent changes in the binaural difference potential in cat auditory brainstem response: Implications about the origin of the binaural interaction component. Hear Res 106:66–82.

Van Hof-Van Duin J (1976) Early and permanent effects of monocular deprivation on pattern discrimination and visuomotor behavior in cats. Brain Res 111:261–276.

Vees AM, Micheva KD, Beaulieu C, Descarries L (1999) Increased number and size of dendritic spines in ipsilateral barrel field cortex following unilateral whisker trimming in postnatal rat. J Comp Neurol 400:110–124.

Wada S-I, Starr A (1983a) Generation of auditory brain stem responses (ABRs). I. Effects of injection of a local anesthetic (procaine HCl) into the trapezoid body of guinea pigs and cat. Electroencephalogr Clin Neurophysiol 56:326–339.

Wada S-I, Starr A (1983b) Generation of auditory brain stem responses (ABRs). II. Effects of surgical section of the trapezoid body on the ABR in guinea pigs and cat. Clin Neurophysiol 56:340–351.

Wada S-I, Starr A (1983c) Generation of auditory brain stem responses (ABRs). III. Effects of lesions of the superior olive, lateral lemniscus and inferior colliculus on the ABR in guinea pig. Electroencephalogr Clin Neurophysiol 56:352–366.

Wada S-I, Starr A (1989) Anatomical bases of binaural interaction in auditory brain-stem responses from guinea pig. Electroencephalogr Clin Neurophysiol 72:535–544.

Whiting B, Moiseff A, Rubio ME (2009) Cochlear nucleus neurons redistribute synaptic AMPA and glycine receptors in response to monaural conductive hearing loss. Neuroscience 163:1264–1276.

Whitton JP, Polley DB (2011) Evaluating the Perceptual and Pathophysiological Consequences of Auditory Deprivation in Early Postnatal Life: A Comparison of Basic and Clinical Studies. J Assoc Res Otolaryngol 12:535–547.

Wiesel TN, Hubel DH (1963) Single-cell responses in striate cortex of kittens deprived of vision in one eye. J Neurophysiol 26:1003–1017.

Xu L, Jen PH-S (2001) The effect of monaural middle ear destruction on postnatal development of auditory response properties of mouse inferior collicular neurons. Hear Res 159:1–13.

Zhou Y, Lai B, Gan WB (2017) Monocular deprivation induces dendritic spine elimination in the developing mouse visual cortex. Sci Rep 7:1–11.

